# Introducing PHJ Media: A Unique Machine Learning -Driven Basal Formulation to Overcome Recalcitrance for Multi-Genotype Micropropagation of *Cannabis sativa* L

**DOI:** 10.64898/2026.07.14.738465

**Authors:** Marco Pepe, Mohsen Hesami, Andrew Maxwell Phineas Jones

## Abstract

Applications of tissue culture are critical for Cannabis sativa L. (cannabis), supporting clonal propagation, germplasm preservation, pathogen elimination, among other biotechnological applications. However, extensive genetic diversity associated with cannabis results in highly variable responses to in vitro conditioning, and no consensus basal media formulation exists to support reproducible micropropagation across genotypes. To address these limitations, a hybridized ensemble-NSGA-II approach was employed for concurrent optimization of individual media components to create a species specific, cultivar inclusive basal salt formulation for cannabis micropropagation. The resulting PHJ media represents a unique formulation that overcomes recalcitrance across a wide array of cannabis cultivars, facilitating improved growth and uniformity for the nine cultivars used in its development and validation. These results remain consistent from explant initiation through multiple rounds of subculture. The ability of PHJ to overcome genotypic recalcitrance is telling of its potential applicability with an array of plant species beyond cannabis. Additionally, robust performance both with and without plant growth regulators underscores the plausible use of PHJ for diverse applications beyond standard micropropagation. Ultimately, this cultivar-inclusive basal medium demonstrates utility for both scientific research and industrial-scale operations.

## 1. Introduction

Tissue culture techniques are quickly becoming standard approaches for clean plant programs, breeding (Hesami et al., 2021a), and clonal mass-production (Shi et al., 2024) of *Cannabis sativa* L. (cannabis). The multitude of existing cannabis genotypes (cultivars) and their unpredictable responses to *in vitro* conditions hinder the standardization of universally applicable best practices. Of particular importance is the basal salt composition, which is the sole source of macronutrients, micronutrients, amino acids, and vitamins to support growth and development of emergent tissues/plantlets (Pepe et al., 2025b). An ideal media formulation should contain specific concentrations of mineral elements and additional compounds to support metabolic function (Sudheer et al., 2022) and should be tailored for a particular plant of interest. However, due to the absence of media optimized for specific species, many are cultured on pre-existing media formulated for other plants. In the case of cannabis, most researchers culture plants on MS or DKW media, originally formulated for tobacco and walnut, respectively. While these media can be used to successfully culture cannabis plants, authors have noted various physiological disorders and signs of nutrient imbalances in both cases (Kastelec et al., 2025; Page et al., 2021; Pepe et al., 2025a).

Customization and optimization of a basal salt formulation for a specific species is complex, involving significant time and effort. This process typically entails the use of stock solutions to mix groups of ionic compounds in various combinations (Phillips and Garda, 2019) and subsequent assessment of efficacy using the One-Factor-at-a-Time (OFAT) approach (Hashizume et al., 2023). While ionic salts represent the means of adding different macro- and micro-nutrients (i.e. KSO_4_), the active factors are the individual ion concentrations (i.e. [K^+^]), which are often donated by multiple salts (Chimdessa, 2020), complicating optimization using the OFAT approach. More importantly, due to the complex interactions among various nutrients, OFAT optimization can lead to false conclusions and sub-optimal results (Pepe et al., 2025b). While other methods such as response surface designs partially overcome this, they still require large numbers of treatments and struggle to efficiently evaluate a complex mixture such as culture media (Beavin et al., 2021). As a result, salts are often grouped and adjusted together to minimize the number of treatments required, but this reduces the ability to optimize individual ions. To overcome the limitations of conventional media optimization procedures, alternative techniques that leverage computational power have been introduced.

Machine learning (ML) represents a subset of artificial intelligence that addresses many of these challenges and is becoming more widely adopted for the optimization of plant tissue culture applications (García-Pérez et al., 2020; Hesami et al., 2021b, 2019; Pepe et al., 2021a, 2021b). The complex, multivariate process of studying *in vitro* plant nutrition involves precise experimental designs and statistical analyses using multiple concomitant parameters that are often difficult to interpret (García-Pérez et al., 2020). To capture the complex, often nonlinear relationships between macro- and micro-nutrients and plant growth traits, advanced machine learning (ML) models are increasingly applied in plant tissue culture optimization. Models such as Support Vector Regression (SVR), Adaptive Neuro-Fuzzy Inference System (ANFIS), and Generalized Regression Neural Network (GRNN) are particularly effective for multivariate systems where interactions among nutrients influence multiple growth responses. Previous studies have proven these models can handle high-dimensional, nonlinear, and interdependent data, making them suitable for predicting plant responses under varying culture conditions, while requiring far fewer treatments (Bozkurt et al., 2024; Hesami et al., 2020; Jafari and Shahsavar, 2020; Pepe et al., 2021a).

Although individual ML models can predict growth responses, they all have different strengths and weaknesses. Combining multiple models through ensemble approaches can enhance prediction accuracy and robustness by leveraging the complementary strengths of different algorithms. Ensemble modeling reduces the individual model bias and variance, resulting in more reliable predictions (Mohammed and Kora, 2023). This is especially relevant in plant tissue culture, where small variations in nutrient concentrations can produce complex and nonlinear effects on growth.

Optimizing culture conditions involves multiple, potentially conflicting objectives, such as maximizing shoot proliferation while promoting canopy expansion. Multi-objective optimization algorithms, such as the Non-dominated Sorting Genetic Algorithm II (NSGA-II), provide a framework to explore trade-offs between growth parameters efficiently. A hybrid approach that combines ensemble models with such optimization techniques enables identification of nutrient combinations that simultaneously enhance multiple growth traits, streamlining the process of culture medium optimization. This approach can be used to simultaneously optimize combinations of influential factors that result in a balance of multiple traits that, together, contribute to overall plant quality (Pepe et al., 2025b; Zhou et al., 2023). For instance, rather than optimizing canopy surface area alone, potentially at the cost of generating basal callus, this method can be used to enhance canopy surface area along with root length, simultaneously.

In addition to prediction and optimization, sensitivity analysis can help identify the relative importance of different macro- and micro-nutrients on plant growth, identifying the most influential factors for specific traits. This offers mechanistic insight into nutrient effects and helps guide targeted adjustments to culture formulation (Jafari and Daneshvar, 2024; Pepe et al., 2025b). The integration of predictive modeling, optimization, and sensitivity analysis thus supports both accurate forecasting of growth outcomes and understanding of nutrient interactions.

Overall, combining ML models, ensemble prediction, multi-objective optimization, and sensitivity analysis establishes a comprehensive framework for optimizing plant tissue culture conditions. This powerful combination forecasts desirable outcomes by simultaneously processing groups of associated input variables to determine their optimal combinations (Ibrahim et al., 2023). By improving the prediction of growth outcomes through effective nutrient management, the protocol optimization process can be streamlined to maximize plant proliferation while reducing the experimental footprint and resource consumption. The approach has successfully been used to optimize tissue culture media and associated protocols for *Juglans regia* L. (Sadat-Hosseini et al., 2022), *Actinidia arguta* (Hameg et al., 2020), *Passiflora caerulea* (Jafari and Daneshvar, 2023), *Chrysanthemum* spp (Hesami et al., 2019), and *Pyrus communis* (Jamshidi et al., 2019), and represents a viable option for optimizing the cannabis basal salt formula for cross-cultivar use. The widespread standardization of this approach for plant tissue culture optimization has prompted the development of detailed reference guides (Pepe et al., 2025b) and open-source platforms to conduct this research (Bethge et al., 2026).

The current investigation enlists the hybridized ensemble-NSGA-II approach to optimize a species specific, cultivar inclusive basal salt formulation for cannabis micropropagation. The experiments identified two optimized basal salt formulations. When tested in a validation experiment with five cannabis genotypes, one optimized formulation in particular afforded improved and predictable node numbers and canopy surface areas (two important features of micropropagation) compared to other commonly used cannabis media formulations. This media formulation was further tested over multiple subcultures on two genotypes, using different explants, and in tests with and without PhytoAx, a proprietary mixture of plant growth regulators (PGRs). The resulting basal media formulation was developed and tested using nine cannabis cultivars throughout the course of experimentation. Overall, the “Pepe, Hesami, Jones medium” (PHJ) resulted in increased growth responses for all explant types, and improved growth during culture initiation. Furthermore, PHJ outperformed DKW media when combined with PhytoAx, demonstrating improved compatibility with PGRs. Thus, this work represents the most comprehensive cannabis media development to-date, resulting in the first custom developed, species specific, cultivar inclusive cannabis basal media formulation. Ultimately, this formulation will advance research and industrial approaches for various cannabis tissue culture applications.

## 2. Materials and Methods

### 2.1. Machine Learning Process

#### 2.1.1. Generation of Training/Testing Dataset

An initial experiment was conducted by creating 122 different combinations of NH_4_NO_3_, K_2_SO_4_, Ca(NO_3_)_2_, CaCl_2,_ KH_2_PO_4_, MgSO_4_, FeSO_4_ · 7H_2_O, Na_2_EDTA · 2H2O, CuSO_4_ · 5H_2_O, MnSO_4_ H_2_O, Na_2_MoO_4_ · 2H_2_O, Zn(NO_3_)_2_ · 6H_2_O, NiSO_4_ · 6H_2_O, and H_3_BO_3_ (Table S1). Added to these treatments were 100mg/L Myo-Inositol, 2mg/L Glycine, 1mg/L Nicotinic acid, 2mg/L Thiamine, 0.004mg/L CoCl_2_, and 0.1mg/L KI. Additionally, each treatment included 0.6% (w/v) agar (Thermo-Fisher Scientific, Waltham, MA), 3% (w/v) sucrose. The pH was adjusted to 5.7 prior to agar addition and autoclaving. Media treatments were dispensed in GA-7 culture vessels (Magenta LLC, Chicago, IL), with 50mL of media in each vessel and 4 vessels per treatment and autoclaved for 20 minutes at 124°C and 15 psi.

Two cannabis genotypes, RTG-X (Roto) (see [Monthony et al., 2021] for SNP-based fingerprint) (Roto-Gro, Bolton, ON, Canada), and BC Black Cherry (BCBC) (The Flowr Group, Kelowna, BC, Canada) were used as stock plant material for the initial experiment. These genotypes were maintained in growth chambers at approximately 25°C, on DKW medium with vitamins (DKW) (Phytotech, Kansas, USA) with 3% (w/v) sucrose, and 0.6% (w/v) agar (Thermo-Fisher Scientific, Waltham, MA), 0.1% (v/v) PPM (Plant Cell Technology, Washington, DC, USA) and pH of 5.7. Light intensity was maintained at 50 µmol/m^2^/s using full-spectrum “Spectra Blade” LED lights emitting 75% Red (600–700 nm), 19% Green (500–600 nm) and 6% Blue (400–500 nm) (Intravision Group, Oslo, Norway).

Shoot tips between 1-2cm in length were used as explants, taken from stock Roto and BCBC maintenance cultures. A total of four explants of each genotype were cultured in four magenta vessels of each treatment, resulting in sixteen explants of each genotype per treatment. Cultures were randomized and maintained in a growth chamber under the conditions described above. After 6-weeks, experimental cultures were taken for imaging and ImageJ was used to assess growth parameters (Rueden et al., 2017), including shoot length, canopy surface area, and node number from the longest shoot.

#### 2.1.2. Machine Learning Models

Three machine learning (ML) algorithms, including support vector regression (SVR), adaptive neuro-fuzzy inference system (ANFIS), and generalized regression neural network (GRNN), were implemented to model the relationship between mineral composition of the culture medium and morphophysiological traits of cannabis plantlets. The concentrations of the 14 salts (NH₄NO₃, K₂SO₄, Ca(NO₃)₂, CaCl₂, KH₂PO₄, MgSO₄, FeSO₄·7H₂O, Na₂EDTA·2H₂O, CuSO₄·5H₂O, MnSO₄·H₂O, Na₂MoO₄·2H₂O, Zn(NO₃)₂·6H₂O, NiSO₄·6H₂O, and H₃BO₃) were used as input variables, and three growth parameters (shoot length, number of nodes, and canopy surface area) were considered as outputs.

From the initial data pool, 492 samples were taken, of which 80% were randomly selected for training and 20% for testing. Prior to model development, the training data were preprocessed by detecting and replacing outliers, followed by normalization of all input features to ensure consistent scaling across variables. The goal of the modeling was to learn a nonlinear mapping *f*: ℝ^14^ → ℝ that predicts each morphological trait from the chemical composition of the medium.

Model performance was evaluated on the independent test set using the coefficient of determination (R²), root mean square error (RMSE), and mean absolute error (MAE), which are defined as:

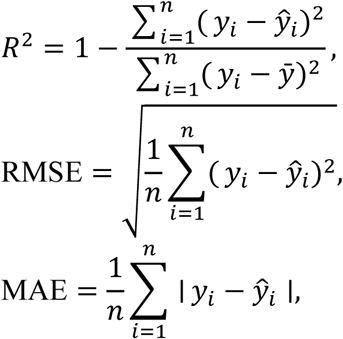

where *y_i_* and *ŷ* are the observed and predicted values, respectively, *ȳ* is the mean of observed values, and *n*is the number of samples. R² measures the proportion of variance explained by the model, RMSE quantifies the average magnitude of prediction errors, and MAE indicates the average absolute deviation of predictions from observations (Chicco et al., 2021).

##### 2.1.2.1. Support Vector Regression (SVR)

Support Vector Regression (SVR) was employed to model the nonlinear relationship between culture medium composition and individual cannabis plantlet growth traits. Separate SVR models were trained for each output variable (shoot length, number of nodes, and canopy surface area). The SVR regression function is defined as:

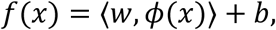

where *ϕ*(*x*) is a nonlinear transformation mapping the input vector *x*into a high-dimensional feature space, *w* is the weight vector, and *b* is the bias. The optimization problem minimizes the regularized ε-insensitive loss:

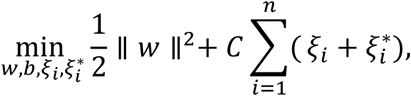

subject to

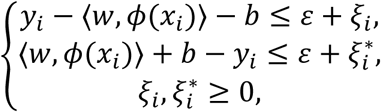

where *C* is the regularization parameter controlling model complexity, and *ε*defines the margin of tolerance for prediction errors. The Radial Basis Function (RBF) kernel was used:

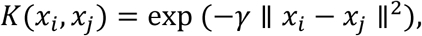

with *γ* controlling the kernel width (Smola et al., 2004).

Hyperparameters *C*, *ε*, and *γ*were optimized separately for each output variable using a grid search. The search ranges were:

1. *C* ∈ [0.1,1000],
2. *γ* ∈ [10^−4^, 10],
3. *ε* ∈ [10^−4^, 1].

Ten-fold cross-validation was performed on the training set to select the hyperparameters that minimized RMSE. SVR models were implemented in MATLAB using the FIRSTVM function. Outlier detection and replacement were applied only on the training set prior to cross-validation to avoid information leakage.

##### 2.1.2.2. Adaptive Neuro-Fuzzy Inference System (ANFIS)

ANFIS was employed to model the nonlinear relationship between culture medium composition and individual cannabis plantlet growth traits. Separate ANFIS models were trained for each output variable (shoot length, number of nodes, and canopy surface area).

ANFIS combines fuzzy logic and artificial neural networks, providing interpretability through fuzzy rules and adaptability through neural learning. Each input variable (e.g., NH₄NO₃ or Ca(NO₃)₂ concentration) was associated with Gaussian membership functions (MFs) representing fuzzy sets such as low and high. A first-order Sugeno fuzzy inference model was used, with rules of the form:

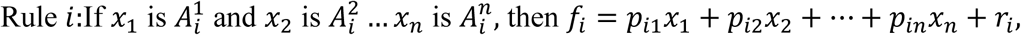

where 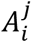 is the fuzzy set of input *x*_j_, and *p*_ij_, *r*_i_ are the linear consequent parameters. The firing strength of the *i*^th^ rule was calculated as:

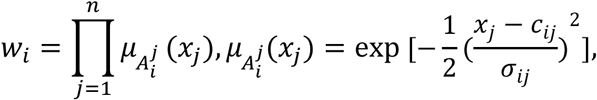

where *c_ij_* and *σ_ij_* are the center and width of the Gaussian MF. Normalized firing strengths were computed as 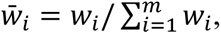, and the final output was obtained as:

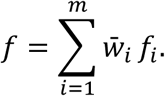

The number of fuzzy rules (*m*) and MFs per input were determined empirically to balance prediction accuracy and computational feasibility. Two MFs per input (28 fuzzy sets total) were used to limit model complexity for the 14-dimensional input space. ANFIS models were trained using a hybrid learning algorithm that combines least-squares estimation for consequent parameters and backpropagation for premise parameters. Training continued until either the RMSE improvement between consecutive epochs was <10⁻⁵ or a maximum of 100 epochs was reached.

Outlier detection and replacement were performed only on the training set prior to model training to avoid information leakage. All ANFIS models were implemented in MATLAB using the ANFIS function.

##### 2.1.2.3. Generalized Regression Neural Network (GRNN)

GRNN was employed to model the nonlinear relationships between culture medium composition and individual cannabis plantlet growth traits. Separate GRNN models were trained for each output variable (shoot length, number of nodes, and canopy surface area). GRNN is a nonparametric regression technique based on kernel density estimation, providing smooth nonlinear function approximation with minimal training time. It was selected for its robustness with small datasets and high-dimensional inputs (Specht, 1991).

The predicted output for a new input *x* is calculated as:

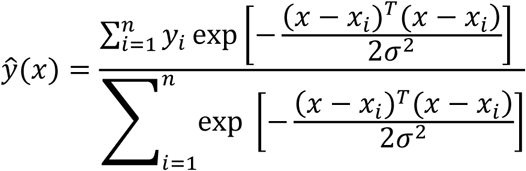

where *x_i_* and *y_i_* are the training samples, and *σ* is the smoothing parameter controlling the spread of the Gaussian kernel. Smaller values of *σ* increase sensitivity to noise, while larger values oversmooth predictions.

The optimal *σ*was determined separately for each output variable via grid search (*σ* ∈ [0.01,1.0]) using ten-fold cross-validation on the training set to minimize RMSE. Outlier detection and replacement were performed only on the training set prior to model training to avoid information leakage.

GRNN models were implemented in MATLAB using a custom-coded network structure with 14 input nodes, one pattern neuron per training sample, and a single output node per growth trait.

#### 2.1.3. Ensemble Model Construction

To improve predictive accuracy and robustness, an ensemble model was constructed by combining the three base learners (SVR, ANFIS, and GRNN). While each base model captures nonlinear relationships independently, their generalization ability may be limited by individual biases and variance.

##### 2.1.3.1. Bagging framework

The ensemble employed a bootstrap aggregating (bagging) approach. Each base model was trained on multiple bootstrap samples (random resampling with replacement) of the training set. For a given input vector *x*, the ensemble output was calculated as:

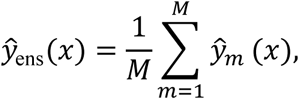

where *ŷ_m_*(*x*) is the prediction of the *m*^th^ base model, and *M* = 3 corresponds to the three trained base learners. Equal weighting was used because preliminary testing showed no significant performance improvement from weighted averaging.

##### 2.1.3.2. Implementation and parameterization

Each base model was trained independently using the optimal hyperparameters described in Sections 2.2.1–2.2.3. Fifty bootstrap replicates of the training set were generated, and ensemble predictions were averaged across replicates to ensure stability. Separate ensembles were constructed for each output variable (shoot length, number of nodes, and canopy surface area). The resulting ensemble model, which demonstrated the highest predictive accuracy, was selected for integration with the NSGA-II algorithm to optimize culture medium composition.

#### 2.1.4. Multi-objective optimization using NSGA-II

To determine the optimal concentrations of macro- and micro-nutrients for maximizing cannabis shoot proliferation, the Non-dominated Sorting Genetic Algorithm II (NSGA-II) was employed. NSGA-II is an evolutionary algorithm designed for multi-objective optimization, simultaneously optimizing conflicting objectives while maintaining population diversity through elitism and crowding distance mechanisms.

In this study, the trained ensemble model (developed from GRNN, SVR, and ANFIS predictors) served as a surrogate fitness function for NSGA-II. The objective was to maximize two growth parameters (i.e., number of nodes and canopy surface area) by optimizing the 14 mineral nutrient inputs: NH₄NO₃, K₂SO₄, Ca(NO₃)₂, CaCl₂, KH₂PO₄, MgSO₄, FeSO₄·7H₂O, Na₂EDTA·2H₂O, CuSO₄·5H₂O, MnSO₄·H₂O, Na₂MoO₄·2H₂O, Zn(NO₃)₂·6H₂O, NiSO₄·6H₂O, and H₃BO₃.

The algorithm was initialized with a population of 200 individuals, each representing a possible combination of the 14 nutrient concentrations within experimentally defined limits. The population was evolved over 1000 generations using the key genetic operators: selection, crossover, and mutation. Roulette wheel selection was used to preferentially propagate higher-performing solutions. Crossover was applied at a rate of 0.7 to recombine parent solutions, while mutation was applied at a rate of 0.5 to maintain diversity and prevent premature convergence.

Non-dominated sorting ranked solutions based on Pareto dominance, and crowding distance preserved diversity along the Pareto front. The algorithm produced a set of Pareto-optimal nutrient combinations, representing trade-offs between the two objectives. The final solution (i.e., the ideal Pareto point) was selected according to the study goal of simultaneously maximizing the number of nodes and canopy surface area. This solution was proposed as the optimal culture medium formulation to enhance cannabis micropropagation efficiency.

#### 2.1.5. Sensitivity Analysis

Sensitivity analysis was conducted to quantify the relative importance of the 14 mineral nutrients (NH₄NO₃, K₂SO₄, Ca(NO₃)₂, CaCl₂, KH₂PO₄, MgSO₄, FeSO₄·7H₂O, Na₂EDTA·2H₂O, CuSO₄·5H₂O, MnSO₄·H₂O, Na₂MoO₄·2H₂O, Zn(NO₃)₂·6H₂O, NiSO₄·6H₂O, and H₃BO₃) on the three growth responses: shoot length, number of nodes, and projected canopy surface area.

The sensitivity of each input variable was assessed using the Variable Sensitivity Error (VSE), which quantifies the performance degradation of the ensemble model when a specific input variable is excluded. For the *j*^th^ input variable, VSE was calculated as:

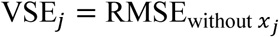

where 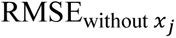 is the root mean square error of the ensemble model when the *j*^th^ variable is omitted from the input set.

The Variable Sensitivity Ratio (VSR) was then defined as the ratio of the VSE to the baseline RMSE of the ensemble model when all input variables were included:

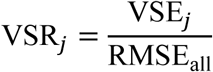

where RMSE_all_ is the model error with all input variables present. A higher VSR indicates greater importance of the corresponding input variable in predicting the output. Variables with VSR > 1 were considered to have a significant influence on the target growth parameters.

### 2.2. Validation Experiments

#### 2.2.1. Validation Experiment 1

Machine learning output predicted sixteen optimized basal media treatments that were tested (Table S2) amongst each other along with DKW and MS with vitamins (MS), and a previously modified MS medium optimized for cannabis (mMS) (Zarei et al., 2023) as control treatments. Aside from the basal formulation, all additional media components remained constant among the treatments, including 3% (w/v) sucrose, and 0.6% (w/v) agar, 0.1% (v/v) PPM, and pH of 5.7. Using the methods from the initial experiment, shoot tip explants from 1-2cm in length from stock Roto cultures maintained on DKW were used and sub-cultured to Magenta GA-7 culture vessels. A total of four explants were cultured per vessel, and each vessel contained 50mL of media. Plants were maintained for 6-weeks under the conditions described in the initial experiment. Plants were then taken for ImageJ analysis (Rueden et al., 2017) to compare shoot length, canopy surface areas, and number of nodes from the longest shoot. Data for this experiment is not shown, as it was used only as a platform to refine treatments for validation experiment 2. Variance components were estimated using restricted maximum likelihood (REML). Degrees of freedom were calculated using Kenward-Roger approximation. Pairwise comparisons were conducted using Tukey’s post hoc analysis at 95% confidence.

#### 2.2.2. Validation Experiment 2

Of the sixteen predicted optimized media treatments from the previous validation experiment, six were chosen for a second round of validation and an additional treatment was included (Table S3) for testing on additional cannabis genotypes. Additional media components included 3% (w/v) sucrose, and 0.6% (w/v) agar, 0.1% (v/v) PPM. and pH of 5.7 for all treatments. These predicted optimized treatments were compared among themselves, and the top two performing treatments were then compared to DKW, MS, and mMS (Zarei et al., 2023). Tested genotypes included Gorilla Glue #4 (GG), Inzane in the Membrane (Inz), Super Sherb (SS) (Dycar Pharmaceuticals, Cranbrook, BC, Canada), White Widow (WW) (Dutch Passion, Amsterdam, the Netherlands), and Roto. These genotypes were being maintained *in vitro* on DKW as described above. Shoot tips 1-2cm long were cultured in Magenta GA-7 vessels containing 50mL of media and maintained for 6-weeks. After the 6-week incubation period, experimental cultures were harvested to compare number of nodes from the longest shoot and canopy surface areas using ImageJ (Rueden et al., 2017). The top two performing media were termed PHJ and PHJ-1. Variance components were estimated using REML, while degrees of freedom for fixed effects were calculated using Kenward-Roger approximation. When significant effects were detected, pairwise post hoc comparison using Tukey was performed. Values for RSD were calculated using the equation outlined in sub-section 2.2.3.1.

#### 2.2.3. Sub-Culture Experiment

To determine the efficacy of the optimized media formulations following sequential sub-cultures, the top performing treatment from the validation experiments (PHJ) was compared to DKW. For this experiment four treatments included PHJ, DKW, and both media with the addition of PhytoAx at 0.075% (v/v) (PHJ+PA, DKW+PA), a proprietary PGR blend (Product ID:P4001; Phytotech, Lenexa, USA). Additional media components were consistent among treatments, as previously described. Treatments were tested with two cannabis genotypes including Roto, and Pedro’s Sweet Sativa (PSS) (CanTx, Puslinch, ON, Canada). Additionally, two explant types were tested, including shoot tips (ST) between 1-2cm and nodal explants (NE) with 2 nodes. For this experiment, four explants of each type (shoot tips and nodal) were cultured in Magenta GA-7 vessels containing 80mL of media, with a total of four vessels per genotype for all treatments. Ultimately, this afforded a total of sixteen explants of each type per genotype, per treatment. For all sub-culture experiments, cultures were maintained under the conditions previously described.

A total of three rounds of sub-culture, including the initial sub-culture and two consecutive rounds, were completed to determine long-term plant performance on different media treatments. After the initial sub-culture from stock plant material, experimental cultures were maintained for 4-weeks before plants were taken for ImageJ analysis and sub-cultured to identical media treatments. These cultures were maintained for 7-weeks under the same conditions before they were taken for ImageJ analysis and an additional round of sub-culture. The final round of sub-culture was maintained for 4-weeks before plants were harvested for ImageJ analysis (Rueden et al., 2017). Both explant types were consistently maintained for all treatments throughout the duration of the experiment. An analysis of each treatment with and without PhytoAx was also completed to determine the efficacy of PhytoAx with different explant types compared to treatments without PhytoAx. Number of shoots, canopy surface area, and number of nodes from longest shoot were quantified using ImageJ analysis. This was done by pooling the data from each genotype with common explant types of the same treatment. Differences among media treatments and differences between explant types were compared using Welch’s t-test.

##### 2.2.3.1. Uniformity of Plants Throughout Subculture

Plantlet trait uniformity throughout subculture was quantified using relative standard deviation (RSD). For shoot number, number of nodes, and canopy surface area, RSD was calculated using the formula:

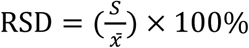

where *S* is the sample standard deviation and *x̅* is the sample mean (Sokal and Rohlf, 2013). For each output variable (shoot number, number of nodes, and canopy surface area), means were calculated by pooling data from both genotypes (Roto and PSS) derived from both explant types (ST and NE). Significance between treatments was compared by conducting an equality of coefficients of variation analysis (Feltz and Miller’s asymptotic test) at 95% confidence using the “cvequality” package in R statistical software version 4.5.0 and RStudio version 2024.12.1 (R Foundation for Statistical Computing, Vienna, Austria).

#### 2.2.4. Culture Initiation Experiment

The effectiveness of media treatments for culture induction was determined by comparing PHJ, DKW with vitamins, both with and without 0.075% (v/v) PhytoAX, for a total of four treatments. Cuttings from two cannabis genotypes, including TRS10 and Super Lemon Haze (SLH), supplied by anonymous donors, were taken from plants grown in a phytotron under conditions describe by Hesami et al. (2023). Fan leaves were removed from cuttings and the bottom tips of cuttings were immediately wrapped in damp paper towel and transported to a laboratory facility for sterilization and culture induction.

For sterilization, cuttings were submerged in cold running water mixed with Sunlight detergent (Henkel Corporation, CT, USA) for 75 minutes. Purox Chlorinating Liquid (Lavo, Montreal, QC, Canada) containing 10.8% (w/w) active sodium hypochlorite was diluted to 30% (v/v) with distilled water. Cuttings were submerged in the dilution and agitated for 15 minutes, then transferred to autoclaved distilled water and agitated. This distilled water washing process was repeated three times for 5 minutes each. Following the sterilization protocol, shoot tips of each genotype between 1-2cm were excised and cultured in Magenta GA-7 vessels containing 80mL of media. A total of 4 explants were cultured per vessel and each treatment was replicated 6 times for a total of 24 explants per genotype, per treatment. For all rounds of the initiation experiments, cultures were maintained under the conditions previously described.

In total, three rounds of culturing, including the initiation culture that followed the sterilization protocol, in addition to two consecutive rounds were completed to compare the efficacy of each treatment for culture induction and maintenance. For the initiation round, cultures were maintained for 5-weeks under the conditions previously described, then plants were sub-cultured to identical media treatments. The newly cultured plants were then maintained for 4-weeks before the next round of sub-culture on identical media. These cultures were then maintained for 4-weeks before they were harvested for ImageJ analysis on the third round of sub-culture. Cultures were assessed after the third round of sub-culture for potential contamination events and to allow the plantlets time to overcome hyperhydricity. Treatments with a minimum of three culture boxes without hyperhydric or contaminated explants were considered for the final analysis, and TRS10 on treatments without PhytoAx were excluded from the final analysis for this reason. Number of nodes on the longest shoot and number of shoots were the parameters measured after three rounds of subculture, for this experiment. Media treatments were compared using Welch’s t-test.

## 3. Results

### 3.1. Predictive Performance of Individual and Ensemble ML Models

The predictive performance of the three individual machine learning models (GRNN, SVR, and ANFIS) and the ensemble model developed using the bagging approach is summarized in Table 1 for shoot length, number of nodes on the longest shoot, and canopy surface area.

**Table 1.**
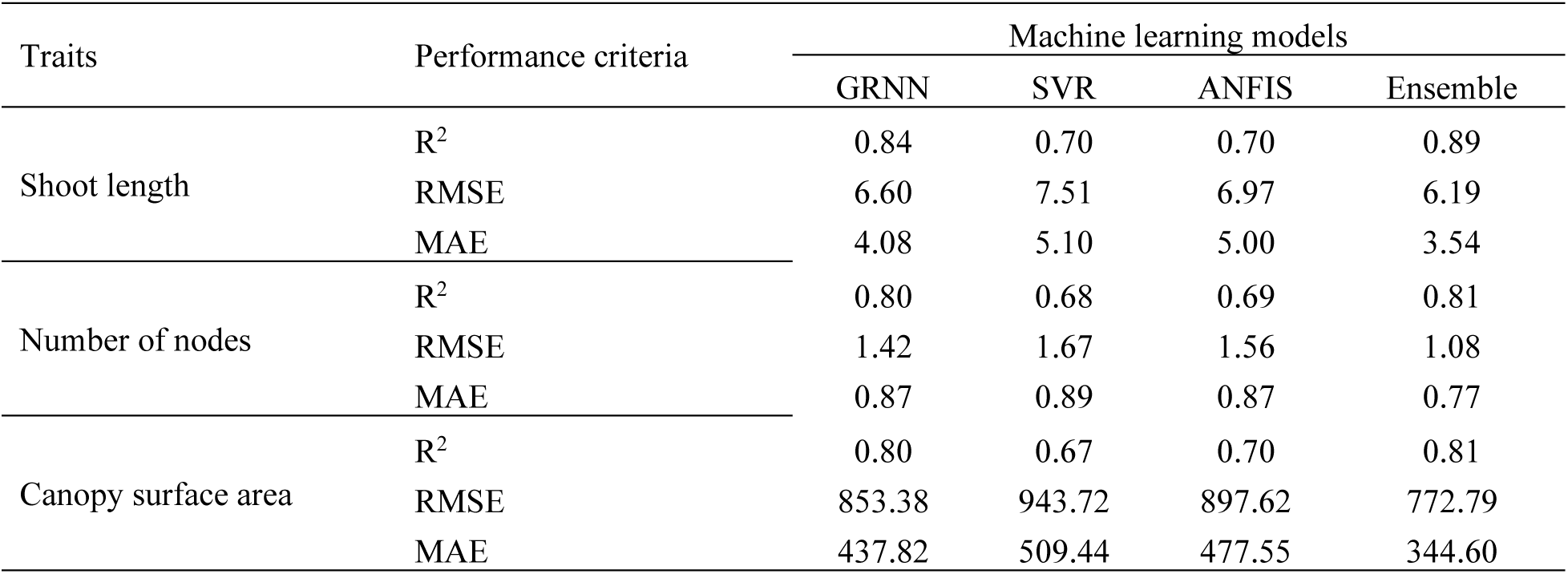
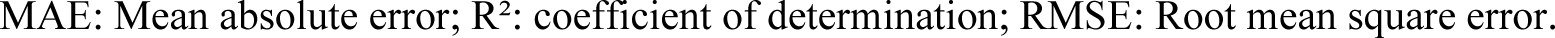
Performance comparison of generalized regression neural network (GRNN), support vector regression (SVR), adaptive neuro-fuzzy inference system (ANFIS), and bagging-based ensemble models for predicting shoot length, number of nodes, and canopy surface area of *Cannabis sativa* L. plantlets based on mineral compositions of the culture medium.

| Traits | Performance criteria | Machine learning models |  |  |  |
| --- | --- | --- | --- | --- | --- |
|  |  | GRNN | SVR | ANFIS | Ensemble |
| Shoot length | R<sup>2</sup> | 0.84 | 0.70 | 0.70 | 0.89 |
|  | RMSE | 6.60 | 7.51 | 6.97 | 6.19 |
|  | MAE | 4.08 | 5.10 | 5.00 | 3.54 |
| Number of nodes | R<sup>2</sup> | 0.80 | 0.68 | 0.69 | 0.81 |
|  | RMSE | 1.42 | 1.67 | 1.56 | 1.08 |
|  | MAE | 0.87 | 0.89 | 0.87 | 0.77 |
| Canopy surface area | R<sup>2</sup> | 0.80 | 0.67 | 0.70 | 0.81 |
|  | RMSE | 853.38 | 943.72 | 897.62 | 772.79 |
|  | MAE | 437.82 | 509.44 | 477.55 | 344.60 |

MAE: Mean absolute error; R<sup>2</sup>: coefficient of determination; RMSE: Root mean square error.

For shoot length, the GRNN model achieved an R² of 0.84, with RMSE and MAE values of 6.60 and 4.08, respectively. The SVR and ANFIS models showed identical R² values of 0.70, with RMSE values of 7.51 and 6.97, and MAE values of 5.10 and 5.00, respectively. The ensemble model yielded an R² of 0.89, with RMSE and MAE values of 6.19 and 3.54, respectively (Table 1).

For the number of nodes, the GRNN model resulted in an R² of 0.80, RMSE of 1.42, and MAE of 0.87. The SVR model produced an R² of 0.68, RMSE of 1.67, and MAE of 0.89, while the ANFIS model showed an R² of 0.69, RMSE of 1.56, and MAE of 0.87. The ensemble model achieved an R² of 0.81, with RMSE and MAE values of 1.08 and 0.77, respectively (Table 1).

For canopy surface area, the GRNN model achieved an R² of 0.80, with RMSE and MAE values of 853.38 and 437.82, respectively. The SVR model resulted in an R² of 0.67, RMSE of 943.72, and MAE of 509.44, while the ANFIS model showed an R² of 0.70, RMSE of 897.62, and MAE of 477.55. The ensemble model produced an R² of 0.81, with RMSE and MAE values of 772.79 and 344.60, respectively (Table 1).

Overall, the ensemble model outperformed the individual models (GRNN, SVR, and ANFIS) across all evaluated traits, as evidenced by higher R² values and lower RMSE and MAE values for shoot length, number of nodes, and canopy surface area (Table 1).

### 3.2. Sensitivity Analysis of Culture Medium Components

The results of the sensitivity analysis, expressed as VSR, for the three output traits are presented in Figure 1.

**Figure 1.**
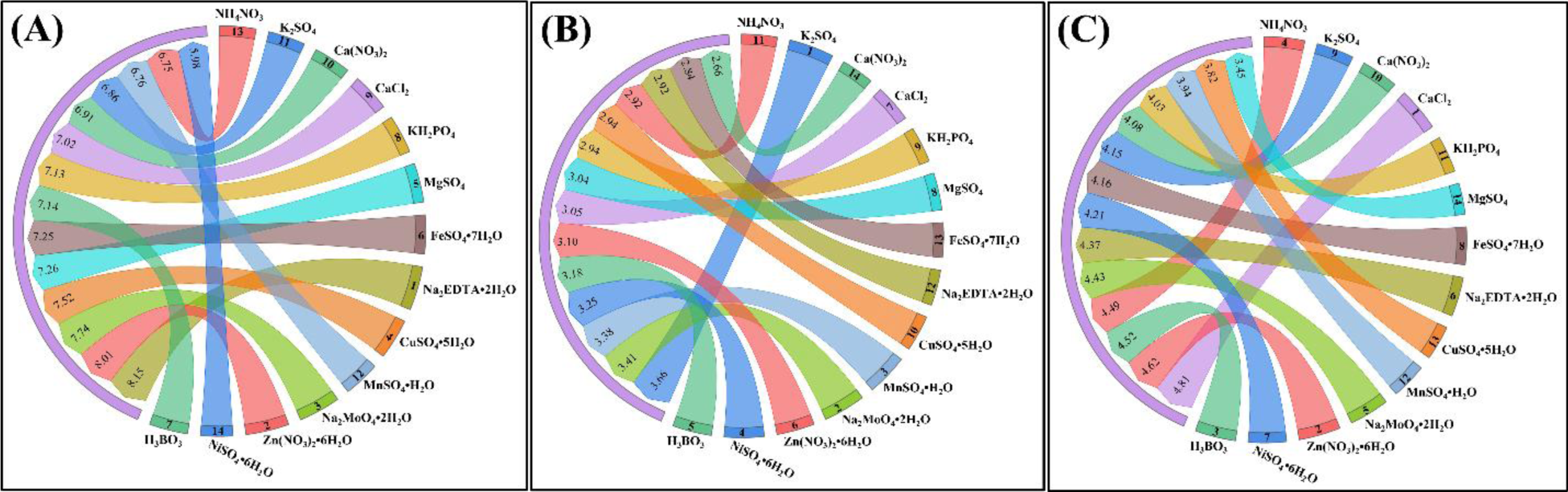
Importance of culture medium components in cannabis micropropagation, as determined by sensitivity analysis of the developed ensemble model. Nutrient contributions are ranked for (A) shoot length, (B) number of nodes, and (C) canopy surface area.

For shoot length (Fig. 1A), VSR values ranged from 5.98 to 8.15 across the evaluated mineral components. The highest VSR was associated with Na₂EDTA·2H₂O (8.15), followed by Zn(NO₃)₂·6H₂O (8.01) and Na₂MoO₄·2H₂O (7.74). Intermediate values were observed for CuSO₄·5H₂O (7.52), MgSO₄ (7.26), FeSO₄·7H₂O (7.25), H₃BO₃ (7.14), KH₂PO₄ (7.13), CaCl₂ (7.02), Ca(NO₃)₂ (6.91), K₂SO₄ (6.86), MnSO₄·H₂O (6.76), and NH₄NO₃ (6.75), while the lowest value was recorded for NiSO₄·6H₂O (5.98).

For the number of nodes (Fig. 1B), VSR values varied between 2.66 and 3.66. The highest VSR was observed for K₂SO₄ (3.66), followed by Na₂MoO₄·2H₂O (3.41) and MnSO₄·H₂O (3.38). Other values included H₃BO₃ (3.18), Zn(NO₃)₂·6H₂O (3.10), CaCl₂ (3.05), MgSO₄ (3.04), KH₂PO₄ (2.94), CuSO₄·5H₂O (2.94), NH₄NO₃ (2.92), Na₂EDTA·2H₂O (2.92), FeSO₄·7H₂O (2.84), and NiSO₄·6H₂O (3.25), with the lowest VSR recorded for Ca(NO₃)₂ (2.66).

For canopy surface area (Fig. 1C), VSR values ranged from 3.45 to 4.81. The highest VSR was associated with CaCl₂ (4.81), followed by Zn(NO₃)₂·6H₂O (4.62), H₃BO₃ (4.52), NH₄NO₃ (4.49), Na₂MoO₄·2H₂O (4.43), Na₂EDTA·2H₂O (4.37), NiSO₄·6H₂O (4.21), FeSO₄·7H₂O (4.16), K₂SO₄ (4.15), Ca(NO₃)₂ (4.08), KH₂PO₄ (4.03), MnSO₄·H₂O (3.94), and CuSO₄·5H₂O (3.82), while the lowest value was observed for MgSO₄ (3.45).

### 3.3. Optimization of Mineral Nutrients Using NSGA-II for Cannabis Micropropagation

The NSGA-II optimization produced two Pareto-optimal nutrient combinations, denoted as PHJ-1 and PHJ, for maximizing the number of nodes and canopy surface area in cannabis shoot proliferation. The concentrations of the 14 mineral nutrients (mg/L) for each predicted optimal point (number of nodes and canopy surface area) are presented in Figure 2.

**Figure 2.**
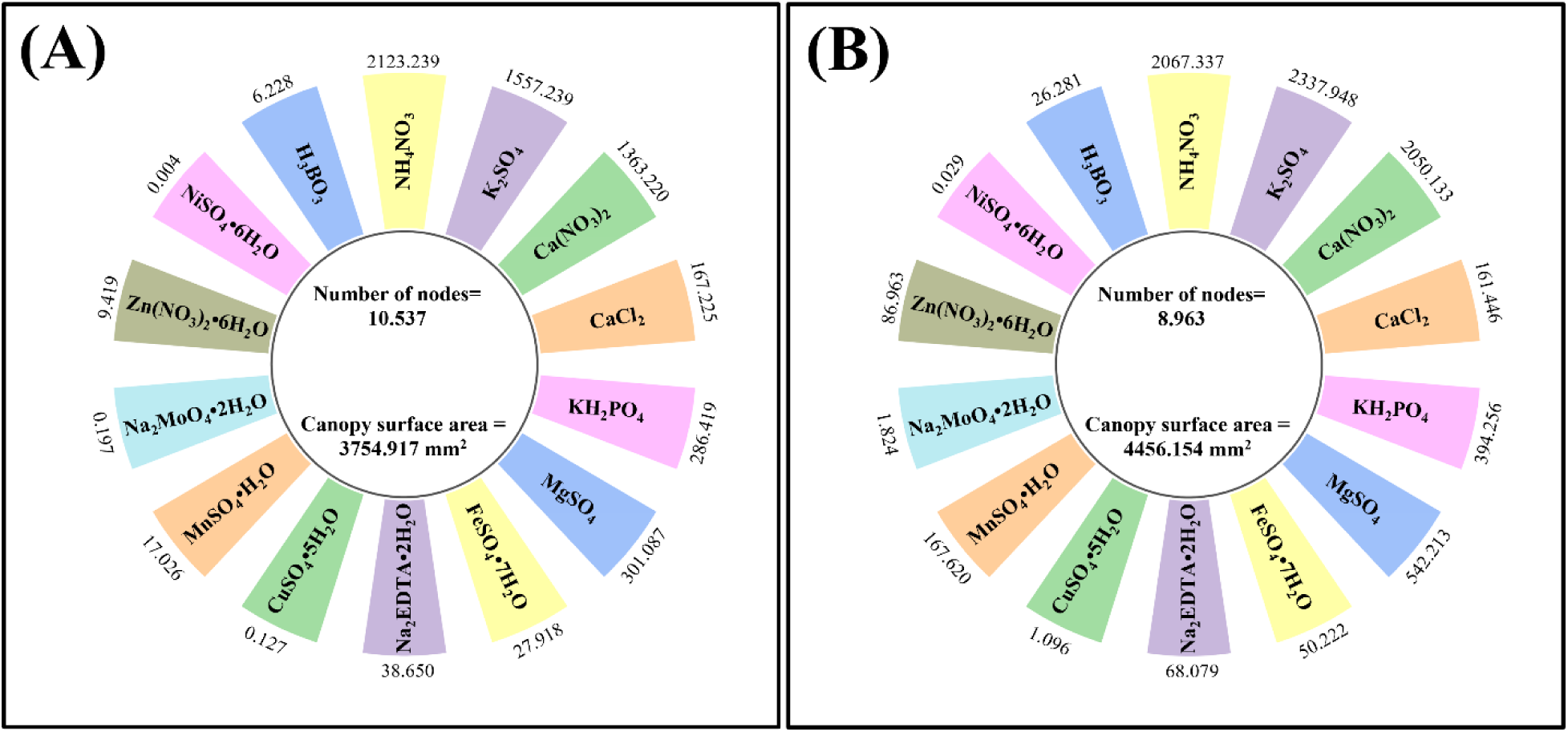
Two Pareto-optimal nutrient combinations, (A) PHJ-1 and (B) PHJ, identified by NSGA-II for maximizing the number of nodes and canopy surface area in micropropagated cannabis shoots. Numbers bordering media components represent concentrations in mg/L. Total additive nutrient salt concentration of PHJ-1 is 5897.998 mg/L. Total additive nutrient salt concentration of PHJ is 7955.447 mg/L.

For PHJ-1 (Fig. 2A), NH₄NO₃ was 2123.24, K₂SO₄ 1557.24, Ca(NO₃)₂ 1363.22, CaCl₂ 167.23, KH₂PO₄ 286.42, MgSO₄ 301.09, FeSO₄·7H₂O 27.92, Na₂EDTA·2H₂O 38.65, CuSO₄·5H₂O 0.13, MnSO₄·H₂O 17.03, Na₂MoO₄·2H₂O 0.20, Zn(NO₃)₂·6H₂O 9.42, NiSO₄·6H₂O 0.004, and H₃BO₃ 6.23.

For PHJ (Fig. 2B), NH₄NO₃ was 2067.34, K₂SO₄ 2337.95, Ca(NO₃)₂ 2050.13, CaCl₂ 161.45, KH₂PO₄ 394.26, MgSO₄ 542.21, FeSO₄·7H₂O 50.22, Na₂EDTA·2H₂O 68.08, CuSO₄·5H₂O 1.10, MnSO₄·H₂O 167.62, Na₂MoO₄·2H₂O 1.82, Zn(NO₃)₂·6H₂O 86.96, NiSO₄·6H₂O 0.03, and H₃BO₃ 26.28.

These values represent the nutrient compositions corresponding to the selected Pareto-optimal solutions for enhancing cannabis micropropagation efficiency.

### 3.4. Validation Experiment 2

Validation experiment 2 focused on the mean comparison of five genotypes on the different media treatments. The rounded averages across genotypes within treatments ± standard errors in mm^2^ are 3638.6 ± 366.17, 2937.1 ± 469.99, 2626.8 ± 281.74, 2373.9 ± 389.03, and 2214.9 ± 261.71 for PHJ, MS, PHJ-1, DKW, and mMS, respectively. In general, PHJ allowed significantly greater canopy surface area than DKW and mMS. No significant differences were shown between mMS, DKW, PHJ-1, or MS. Although PHJ allowed a greater overall mean data cluster of canopy surface area for all the tested genotypes, no significant differences were shown between PHJ, PHJ-1, or MS. Canopy surface area RSD values for PHJ, MS, PHJ-1, DKW, and mMS are 45.00%, 71.56%, 43.05%, 73.29%, and 52.84%, respectively (Fig. 3). Across all media, WW showed the highest mean canopy surface area (4015.56mm^2^), while GG had the lowest mean canopy surface area (1904.83mm^2^).

**Figure 3.**
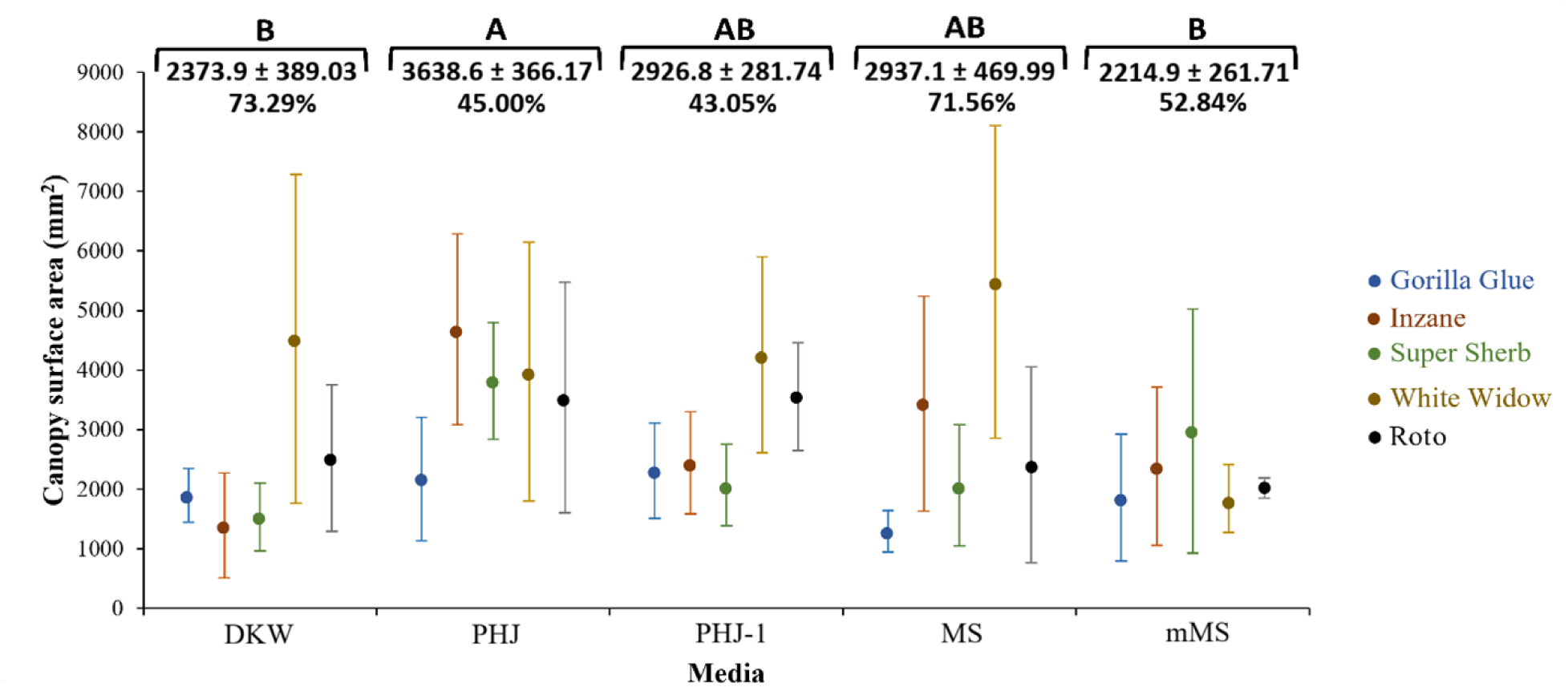
Mean canopy surface area of genotypes tested on different media treatments. Error bars represent standard error of the mean for individual genotypes within treatments. Letters represent significance based on rounded grouped averages across all genotypes within treatments, when compared to those of other treatments with 95% confidence. Grouped averages of genotypes ± standard errors in mm^2^ are shown beneath letters. Relative standard deviations (RSD) are shown under grouped averages.

When comparing mean number of nodes, the rounded averages across genotypes within treatments ± standard errors in mm^2^ are 9 ± 0.3, 8 ± 0.3, 7 ± 0.3, 7 ± 0.3, and 7 ± 0.2 for PHJ, PHJ-1, DKW, MS, and mMS, respectively. Although PHJ allowed the greatest number of nodes overall, there were no significant differences in the number of nodes produced between PHJ and PHJ-1. Both PHJ and PHJ-1 allowed significantly higher node numbers than DKW, MS, and mMS, while there were no significant differences in the number of nodes produces by DKW, MS, or mMS. Number of nodes RSD values for PHJ, PHJ-1, DKW, MS, and mMS, are 29.51%, 28.66%, 34.51%, 37.01%, and 30.08%, respectively (Fig. 4). Across all media, WW averaged the most nodes (9.10), while SS averaged the least number of nodes (6.61).

**Figure 4.**
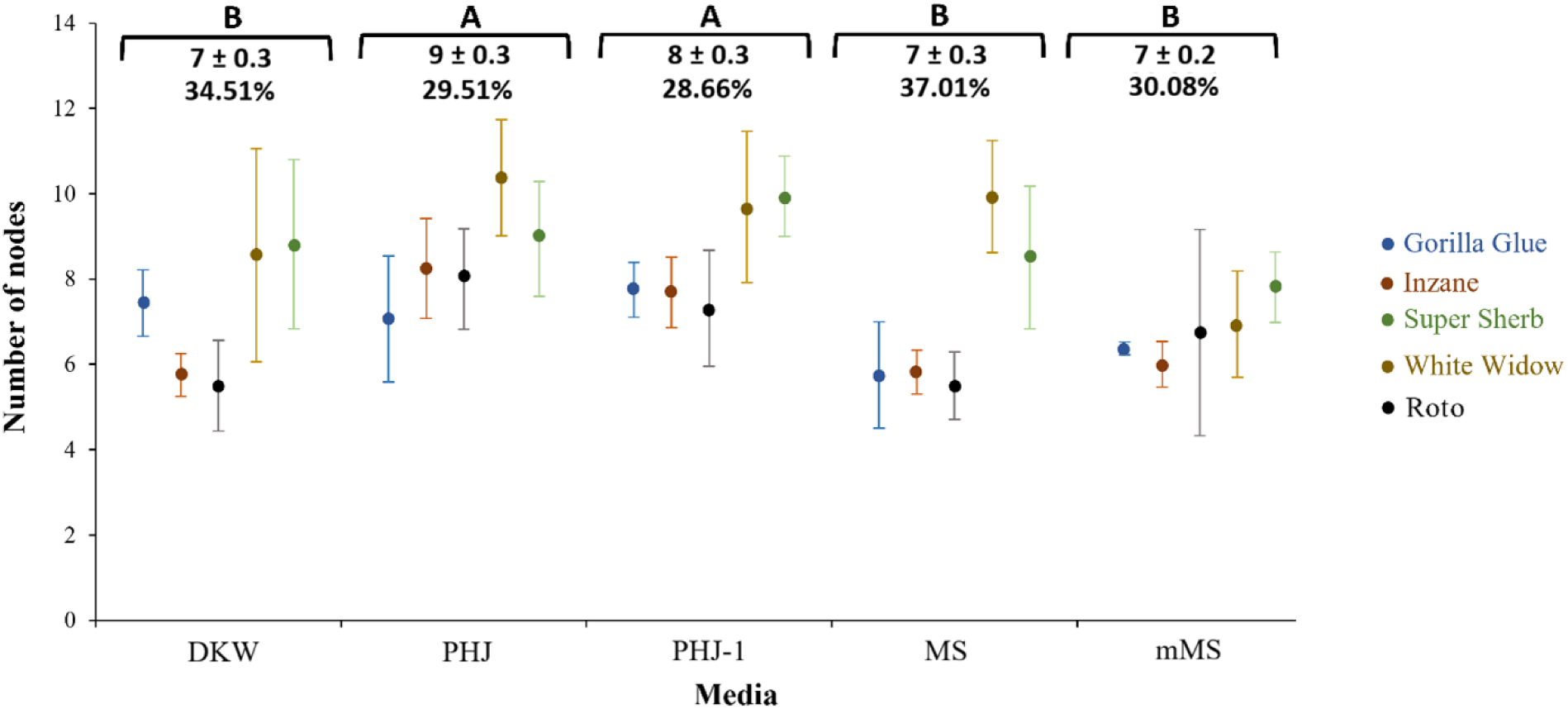
Mean number of nodes produced by genotypes tested on different media treatments. Error bars represent standard error of the mean for individual genotypes within treatments. Letters represent significance based on rounded grouped averages across genotypes within treatments, when compared to those of other treatments with 95% confidence. Letters represent significance based on rounded grouped averages across all genotypes within treatments, when compared to those of other treatments with 95% confidence. Grouped rounded averages of genotypes ± standard errors in mm^2^ are shown beneath letters. Relative standard deviations (RSD) are shown under grouped rounded averages.

### 3.5. Subculture Experiments

#### 3.5.1. Subculture Phase 1

Significantly greater number of shoots were produced on PHJ compared to DKW, except for PSS NC, where no significant differences between PHJ and DKW were measured. Node numbers, in all cases, were greater on PHJ than DKW. For both genotypes, PHJ NC allowed significantly greater canopy surface area compared to DKW NC, while DKW ST resulted in greater canopy surface area than PHJ ST, though these differences were insignificant (Table 2).

**Table 2.** Subculture 1 assessment of media treatments based on explant type, including shoot tip (ST) and nodal cutting (NC). Values represent averages of the parameters tested ± standard error of the mean. Significance, with 95% confidence, is noted with asterisks.

| Parameter | Media | Roto NC | Roto ST | PSS NC | PSS ST |
| --- | --- | --- | --- | --- | --- |
| Shoot Number | PHJ | 2.1 ± 0.12* | 1.7 ± 0.21* | 1.8 ± 0.23 | 2.0 ± 0.10* |
| | DKW | 1.6 ± 0.16 | 1.1 ± 0.07 | 1.8 ± 0.27 | 1.4 ± 0.16 |
| Number of Nodes | PHJ | 7.1 ± 0.30* | 7.8 ± 0.19* | 9.4 ± 0.31** | 9.9 ± 0.48** |
| | DKW | 5.5 ± 0.43 | 6.7 ± 0.39 | 6.1 ± 0.44 | 6.8 ± 0.23 |
| Canopy Area | PHJ | 3441.7 ± 60.88** | 1494.9 ± 89.74 | 2799.6 ± 637.37* | 1635.0 ± 203.35 |
| | DKW | 1618.3 ± 262.26 | 1675.1 ± 174.92 | 1309.2 ± 80.79 | 1936.3 ± 265.51 |

#### 3.5.2. Subculture Phase 2

Shoot number was greater for both genotypes and explant types on PJH, though there were no significant differences. In all cases, number of nodes was greater on PHJ, with the only significant difference occurring with Roto ST. There were no significant differences in canopy surface area. In general, canopy surface area was greater on PHJ compared to DKW, with the exception being PSS ST Table 3.

**Table 3.** Subculture 2 assessment of media treatments based on explant type, including shoot tip (ST) and nodal cutting (NC). Values represent averages of the parameters tested ± standard error of the mean. Significance, with 95% confidence, is noted with asterisks.

| Parameter | Media | Roto NC | Roto ST | PSS NC | PSS ST |
| --- | --- | --- | --- | --- | --- |
| Shoot Number | PHJ | 2.4 ± 0.30 | 1.8 ± 0.50 | 2.1 ± 0.16 | 1.5 ± 0.34 |
| | DKW | 1.9 ± 0.16 | 1.1 ± 0.07 | 1.6 ± 0.22 | 1.4 ± 0.16 |
| Number of Nodes | PHJ | 9.1 ± 0.75 | 9.5 ± 0.20* | 9.7 ± 0.54 | 9.5 ± 0.76 |
| | DKW | 8.4 ± 0.62 | 8.3 ± 0.37 | 6.9 ± 0.91 | 8.8 ± 0.60 |
| Canopy Area | PHJ | 4693.56 ± 611.02 | 1973.46 ± 240.09 | 3113.24 ± 967.36 | 2190.9 ± 554.50 |
| | DKW | 4539.5 ± 916.63 | 1891.4 ± 82.55 | 3023.7 ± 895.20 | 4091.5 ± 991.29 |

#### 3.5.3. Subculture Phase 3

Significantly greater shoot numbers occurred on PHJ for Roto NC and PSS NC compared to DKW. Albeit not statistically significant, greater shoot numbers occurred on PHJ for PSS ST compared to DKW, and DKW for Roto ST compared to PHJ. Greater number of nodes occurred on PHJ, in all cases except for PSS ST. Significantly greater numbers of nodes occurred on PHJ for Roto ST and PSS NC. Greater canopy surface areas occurred on PHJ in all cases, except for PSS ST. Significantly greater canopy surface areas occurred on PHK for Roto ST (Table 4).

**Table 4.** Subculture 3 assessment of media treatments based on explant type, including shoot tip (ST) and nodal cutting (NC). Values represent averages of the parameters tested ± standard error of the mean. Significance, with 95% confidence, is noted with asterisks.

| Parameter | Media | Roto NC | Roto ST | PSS NC | PSS ST |
| --- | --- | --- | --- | --- | --- |
| Shoot Number | PHJ | 2.1 ± 0.33* | 1.0 ± 0.00 | 2.44 ± 0.12** | 1.6 ± 0.19 |
| | DKW | 1.8 ± 0.16 | 1.2 ± 0.12 | 1.4 ± 0.12 | 1.3 ± 0.06 |
| Number of Nodes | PHJ | 6.9 ± 0.56 | 8.4 ± 0.44* | 7.6 ± 0.16** | 8.3 ± 0.66 |
| | DKW | 5.6 ± 0.26 | 5.9 ± 0.53 | 5.6 ± 0.39 | 8.4 ± 0.74 |
| Canopy Area | PHJ | 1927.4 ± 226.11 | 2439.4 ± 175.42* | 1686.9 ± 164.37 | 2005.6 ± 146.20 |
| | DKW | 1778.6 ± 136.94 | 1912.5 ± 172.66 | 1646.9 ± 117.93 | 2324.4 ± 433.16 |

#### 3.5.4. Subculture Phase 1 with PhytoAx

Shoot number, in all cases, were greater on PHJ+PA, though the only significant difference was with Roto NC. Similarly, PHJ+PA allowed greater number of nodes, in all cases, though only significantly different with Roto NC. Canopy surface area on PHJ+PA was significantly greater than DKW+PA for Roto of both explant types. For PSS NC, canopy surface area was greater on DKW+PA, while canopy surface area was greater on PHJ+PA for PSS ST, in both cases, these differences were not statistically significant (Table 5).

**Table 5.** Subculture 1 assessment of media treatments with PhytoAx based on explant type, including shoot tip (ST) and nodal cutting (NC). Values represent averages of the parameters tested ± standard error of the mean. Significance, with 95% confidence, is noted with asterisks.

| Parameter | Media | Roto NC | Roto ST | PSS NC | PSS ST |
| --- | --- | --- | --- | --- | --- |
| Shoot Number | PHJ+PA | 4.0 ± 0.42** | 1.8 ± 0.26 | 4.9 ± 0.77 | 3.6 ± 1.03 |
| | DKW+PA | 1.6 ± 0.07 | 1.4 ± 85.73 | 3.9 ± 0.98 | 2.4 ± 0.51 |
| Number of Nodes | PHJ+PA | 8.4 ± 0.54** | 6.0 ± 0.31 | 10.5 ± 0.91 | 10.5 ± 0.62 |
| | DKW+PA | 5.1 ± 0.48 | 5.4 ± 0.77 | 9.5 ± 0.35 | 8.2 ± 1.00 |
| Canopy Area | PHJ+PA | 2256.4 ± 408.50* | 825.1 ± 96.98* | 5041.1 ± 277.19 | 4003.3 ± 416.54 |
| | DKW+PA | 629.1 ± 114.56 | 476.3 ± 85.73 | 5721.6 ± 747.63 | 2426.9 ± 499.60 |

#### 3.5.5. Subculture Phase 2 with PhytoAx

For both explant types, PHJ+PA produced greater shoot numbers, node numbers, and canopy surface areas than DKW+PA. Significantly higher shoot numbers occurred on PHJ+PA with Roto NC and PSS ST. Node numbers were significantly greater on PHJ+PA with Roto NC and PSS ST. Canopy surface area was significantly greater on PHJ+PA with Roto NC, ST, and PSS, ST (Table 6).

**Table 6.** Subculture 2 assessment of media treatments with PhytoAx based on explant type, including shoot tip (ST) and nodal cutting (NC). Values represent averages of the parameters tested ± standard error of the mean. Significance, with 95% confidence, is noted with asterisks.

| Parameter | Media | Roto NC | Roto ST | PSS NC | PSS ST |
| --- | --- | --- | --- | --- | --- |
| Shoot Number | PHJ+PA | 4.2 ± 0.19*** | 1.8 ± 0.21 | 6.0 ± 0.26 | 7.8 ± 0.44*** |
| | DKW+PA | 1.7 ± 0.21 | 1.6 ± 0.16 | 5.4 ± 0.77 | 2.4 ± 0.06 |
| Number of Nodes | PHJ+PA | 10.5 ± 0.94** | 7.5 ± 0.4 | 11.9 ± 0.53 | 12.2 ± 0.54* |
| | DKW+PA | 5.9 ± 0.22 | 7.3 ± 0.57 | 11.8 ± 0.57 | 9.6 ± 0.31 |
| Canopy Area | PHJ+PA | 4200.5 ± 722.31** | 1444.5 ± 46.35** | 9818.2 ± 1621.33 | 9851.9 ± 1167.2* |
| | DKW+PA | 937.7 ± 466.05 | 1216.4 ± 79.22 | 7046.1 ± 352.98 | 4572.5 ± 798.12 |

#### 3.5.6. Subculture Phase 3 with PhytoAx

Overall, for all parameters measured, genotypes, and explant types, PHJ+PA produced greater growth than DKW+PA. Significantly greater shoot and node numbers occurred on PHJ+PA for all genotypes and explant types compared to DKW+PA. For canopy surface area PSS ST on PHJ+PA was significantly greater than DKW+PA (Table 7).

**Table 7.**
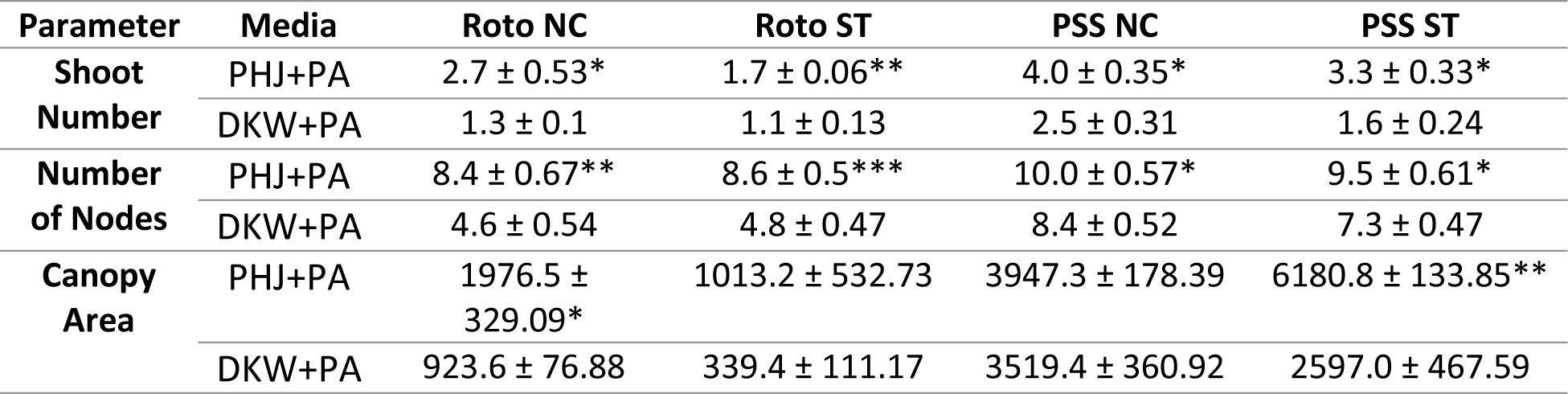
Subculture 3 assessment of media treatments with PhytoAx based on explant type, including shoot tip (ST) and nodal cutting (NC). Values represent averages of the parameters tested ± standard error of the mean. Significance, with 95% confidence, is noted with asterisks.

### 3.6. Notes on Specimen Uniformity and Quality Throughout Subculture

#### 3.6.1. Subculture Phase 1

Following 4-weeks in the subculture 1 phase, more uniform responses were characteristic of specimens grown on PHJ compared to DKW for shoot number based on RSD (PHJ: 18.85%, DKW: 27.71%). For number of nodes, RSD was nearly identical on both media treatments (PHJ: 15.26%, DKW: 13.70%). Canopy surface area RSD of plantlets was greater on PHJ (44.16%) compared to DKW (26.95%). No significant differences were found between treatments, based on RSD following subculture phase 1 (Fig. 5).

**Figure 5.**
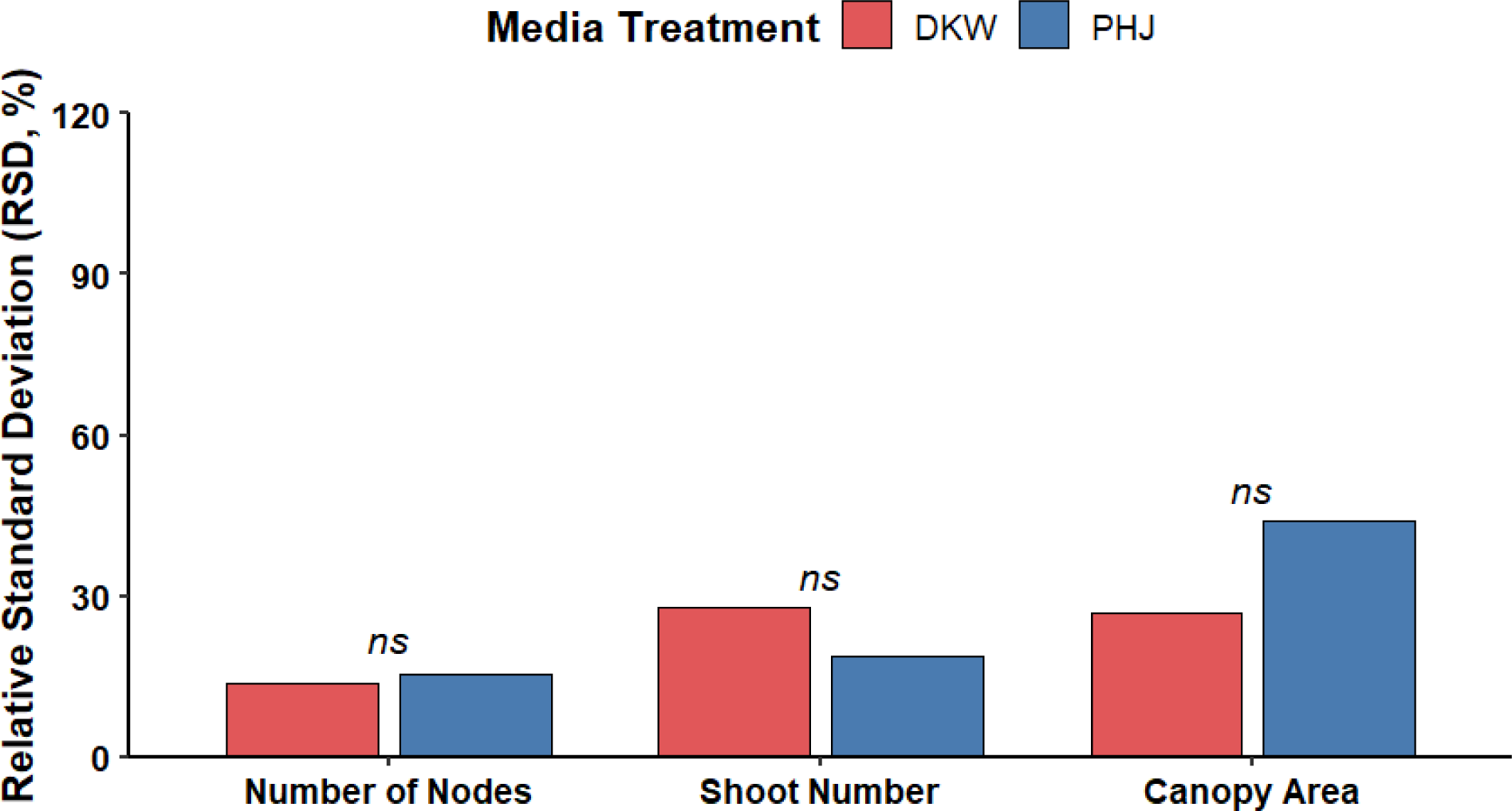
Comparison of overall plantlet uniformity following the subculture 1 phase. The RSD for DKW and PHJ are reported for each response variable quantified. Significance is noted with 95% confidence.

Plantlet sizes and morphologies generally appeared healthier among ST and NC of both genotypes when cultured on PHJ. Single dominant shoots were common from ST. Multiple co-dominant shoots were common from NC on PHJ for each genotype. In general, NC delivered greater canopy surface areas for both genotypes when grown on PHJ, while ST delivered greater canopy surface areas for both genotypes when grown on DKW (Table 2). Overall, PSS responded better to culture conditions than Roto (Fig. 6).

**Figure 6.**
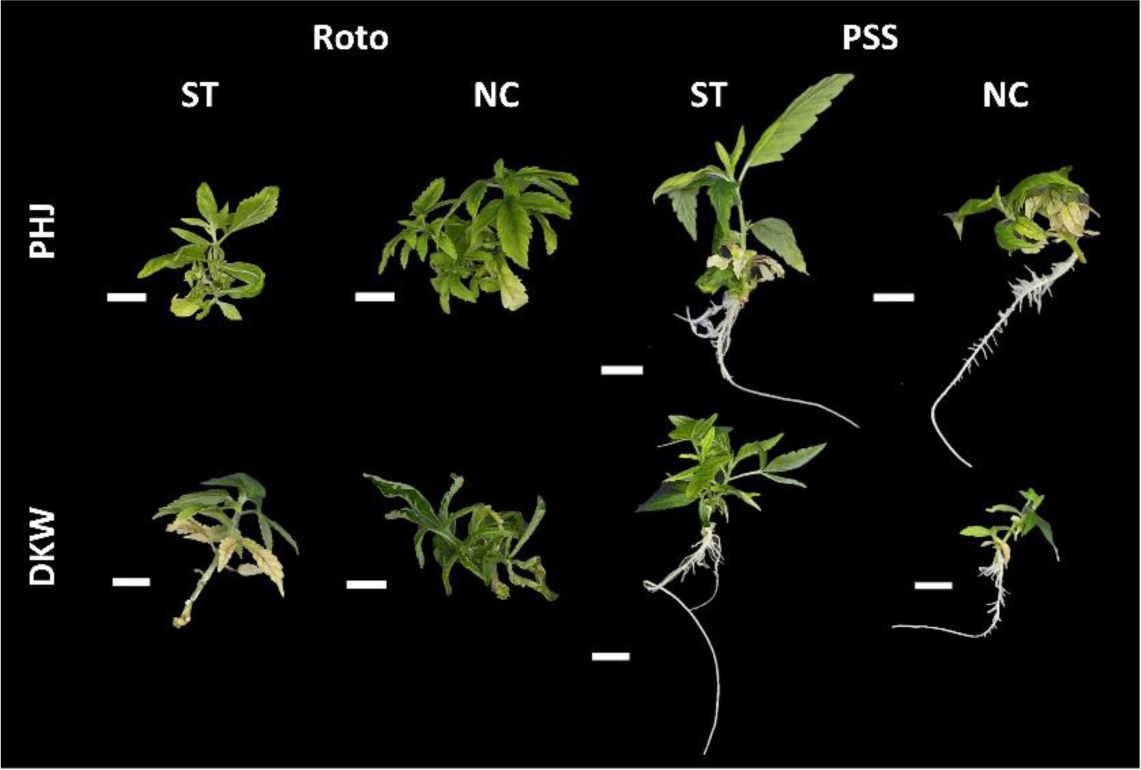
Representative specimens after the 4-week subculture 1 phase. Depicted are genotypes Roto and PSS grown from shoot tips (ST) and nodal cuttings (NC) on PHJ and DKW. Scale bars represent 10mm length.

#### 3.6.2. Subculture Phase 2

Following the 7-weeks phase of subculture 2, RSD indicated that number of nodes was more uniform of plantlets grown on PHJ (11.56%) compared to DKW (16.90%). For canopy surface area, RSD between treatments was nearly identical (PHJ: 53.40, DKW: 53.01). Shoot numbers of plantlets grown on PHJ (36.27%) was higher than DKW (27.24%). Overall, no significant differences were noted for uniformity based on RSD (Fig. 7).

**Figure 7.**
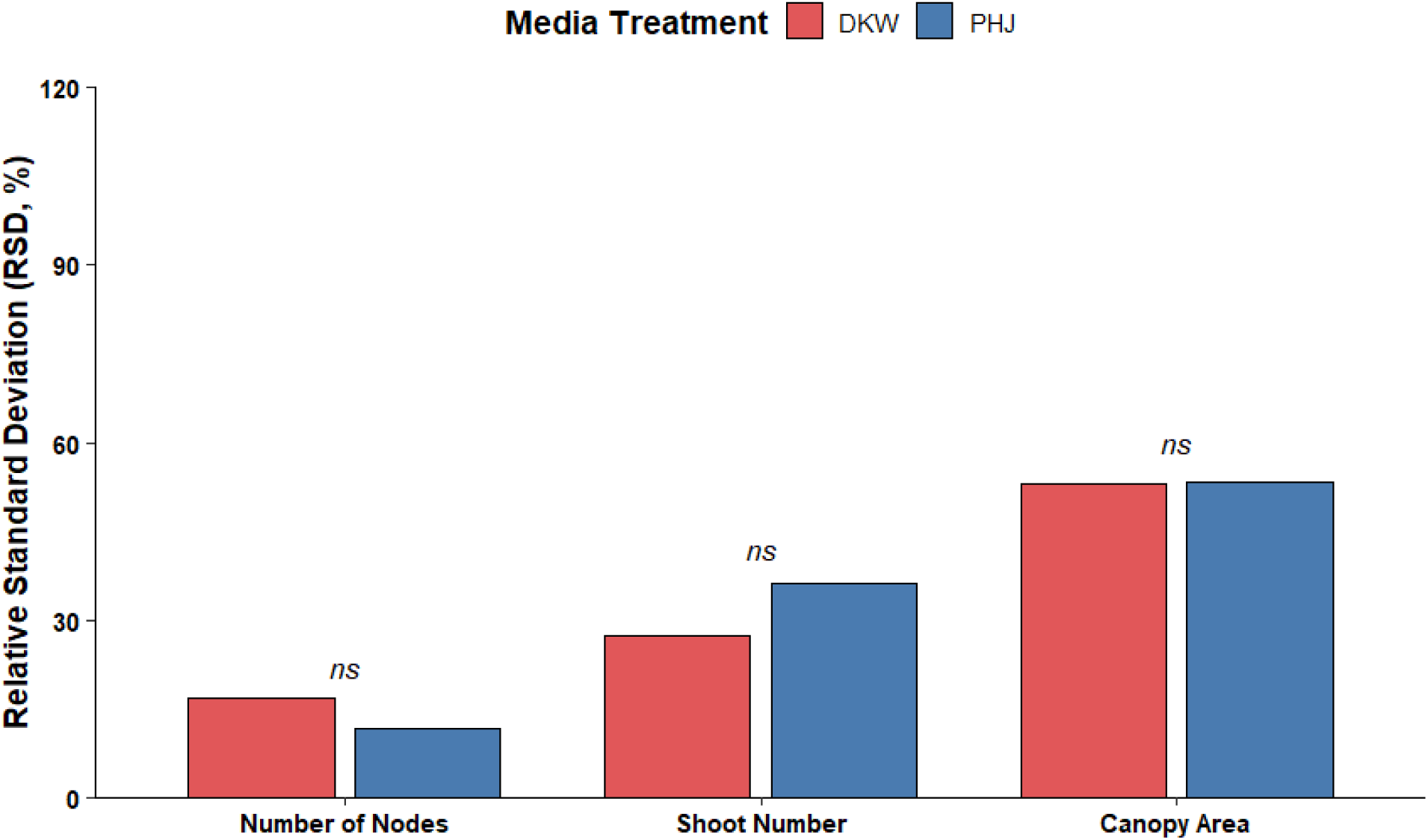
Comparison of overall plantlet uniformity following the subculture 2 phase. The RSD for DKW and PHJ are reported for each response variable quantified. Significance is noted with 95% confidence.

The majority of plantlets of each genotype, explant type, on both PHJ and DKW had rooted following the subculture 2 phase. However, there were several occurrences of basal callus development for Roto ST on DKW. These plantlets did not develop roots. In general, PSS responded better to culture conditions than Roto (Fig. 8).

**Figure 8.**
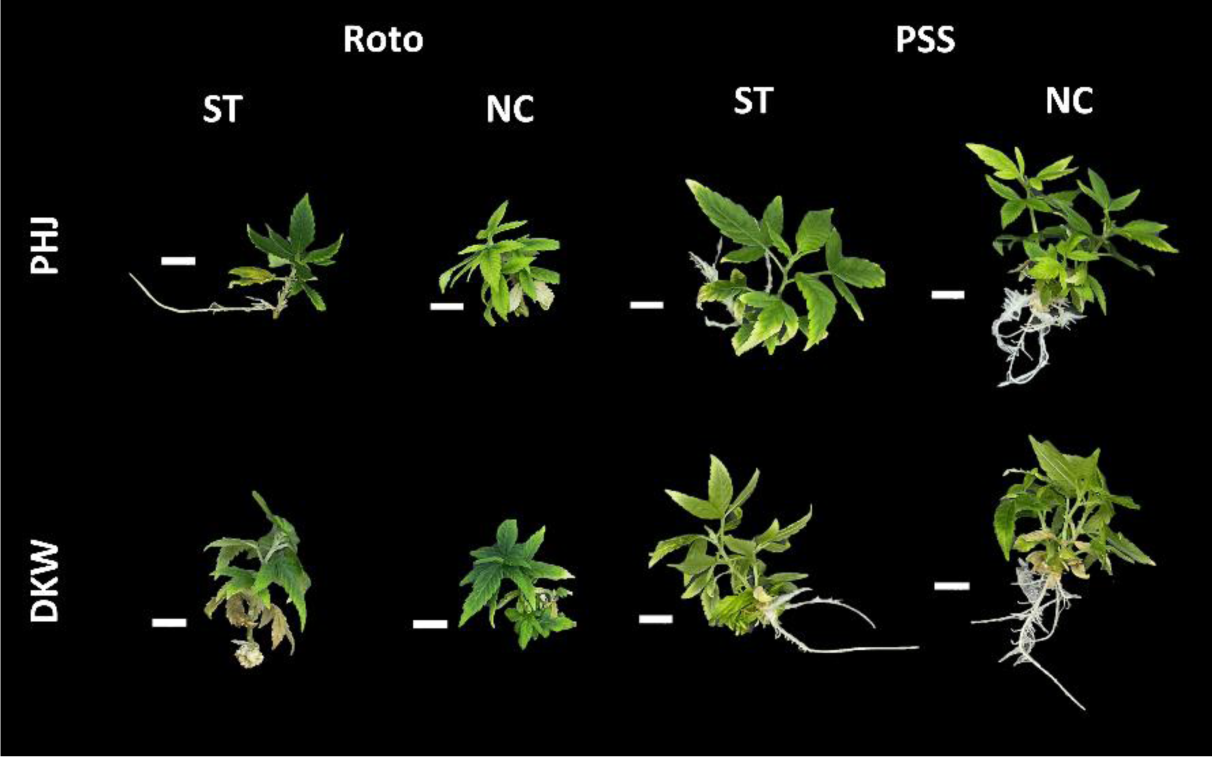
Representative specimens after the 7-week subculture 2 phase. Depicted are genotypes Roto and PSS grown from shoot tips (ST) and nodal cuttings (NC) on PHJ and DKW. Scale bars represent 10mm length.

#### 3.6.3. Subculture Phase 3

The 4-week subculture 3 phase resulted in significantly more uniform node numbers for plantlets grown on PHJ (13.80%) compared to DKW (23.70%) based on RSD. Canopy surface area RSD was lower for plantlets grown on PHJ (22.03%) compared to DKW (27.07%). Shoot number RSD was greater for plants grown on PHJ (37.45%) compared to DKW (22.45%) (Fig. 9).

**Figure 9.**
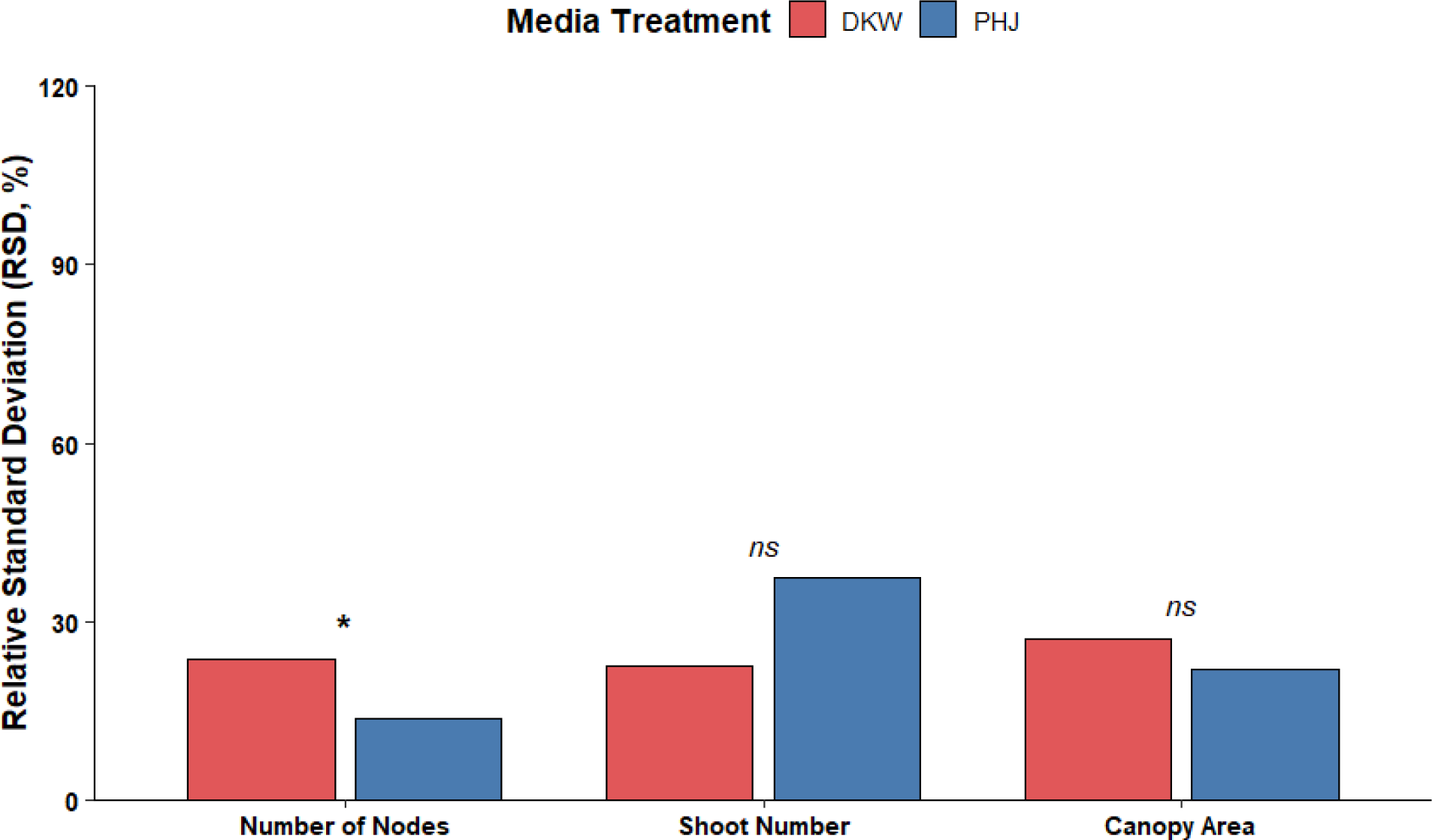
Comparison of overall plantlet uniformity following the subculture 3 phase. The RSD for DKW and PHJ are reported for each response variable quantified. Significance is noted with 95% confidence.

In general, larger plantlets with more shoots and nodes were characteristic of both genotypes and explant types cultured on PHJ compared to DKW (Table 4). As with previous subcultures, PSS responded better to culture conditions than Roto. However, both genotypes responded better to PHJ compared to DKW (Fig. 10).

**Figure 10.**
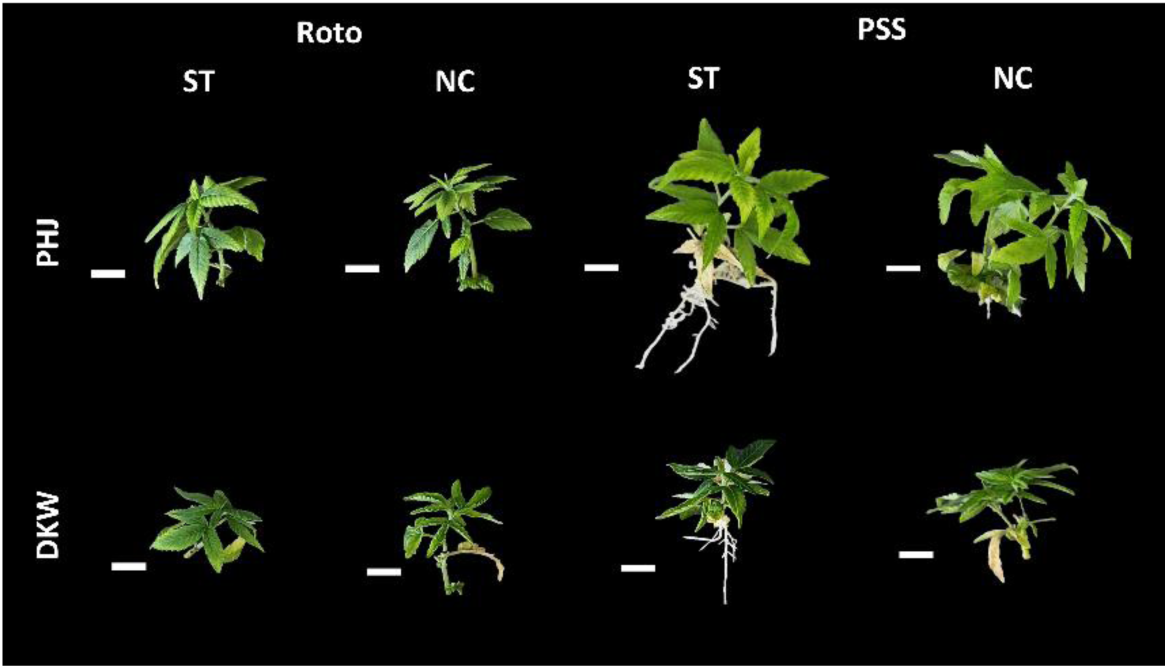
Representative specimens after the 4-week subculture 3 phase. Depicted are genotypes Roto and PSS grown from shoot tips (ST) and nodal cuttings (NC) on PHJ and DKW. Scale bars represent 10mm length.

#### 3.6.4. Characterization of the “PhytoAx Phenotype”

Throughout experimentation, treatments that included PhytoAx tended to develop a distinct phenotype, in which we identify as the “PhytoAx phenotype”. The PhytoAx phenotype is characterized by the occurrence of basal callus, long internodes, and spindly or curled, and sometimes chlorotic leaves (Fig. 11). This phenotype was most prominent with explants obtained from subculture 2 and was more pronounced on DKW+PA compared to PHJ+PA treatments. Generally, the emergence of the PhytoAx phenotype began by the end of the subculture 1 phase with the occurrence of swollen basal stems and chlorotic leaves (Fig. 13). By the end of the subculture 2 phase, the swollen basal stems developed basal callus, leaves became curled and thin, and shoots would multiply and elongate (Fig. 15). By the end of the subculture 3 phase, most plants had reverted to a more normal phenotype, with more regular leaf size, shape, and colour, and fewer elongated stems, but would retain basal callus, and multiple shoots (Fig. 17).

**Figure 11.**
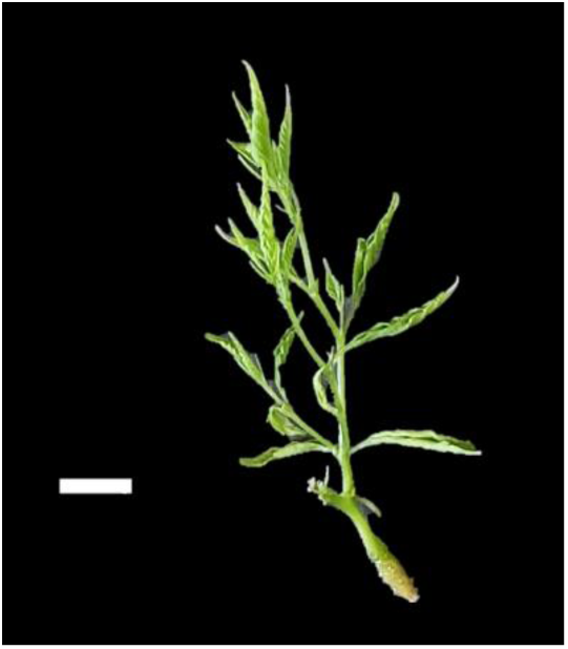
Example of specimen plant displaying the PhytoAx phenotype, characterized by the occurrence of basal callus, numerous shoot tips, long internodes, and spindly or curled, and sometimes chlorotic leaves. Depicted is PSS on DKW+PA NC, obtained after subculture phase 3. Scale bar represents 10mm length.

#### 3.6.5. Subculture Phase 1 with PhytoAx

After 4-weeks in the subculture 1 phase, RSD trends showed greater uniformity of plantlets grown on PHJ+PA. Number of nodes, shoot number, and canopy surface area of planets grown on PHJ+PA (25.09%, 47.31%, and 58.20%, respectively) were all numerically more uniform than those grown on DKW+PA (32.74%, 60.58%, and 100.64%). However, the differences in relative uniformity were insignificant (Fig. 12).

**Figure 12.**
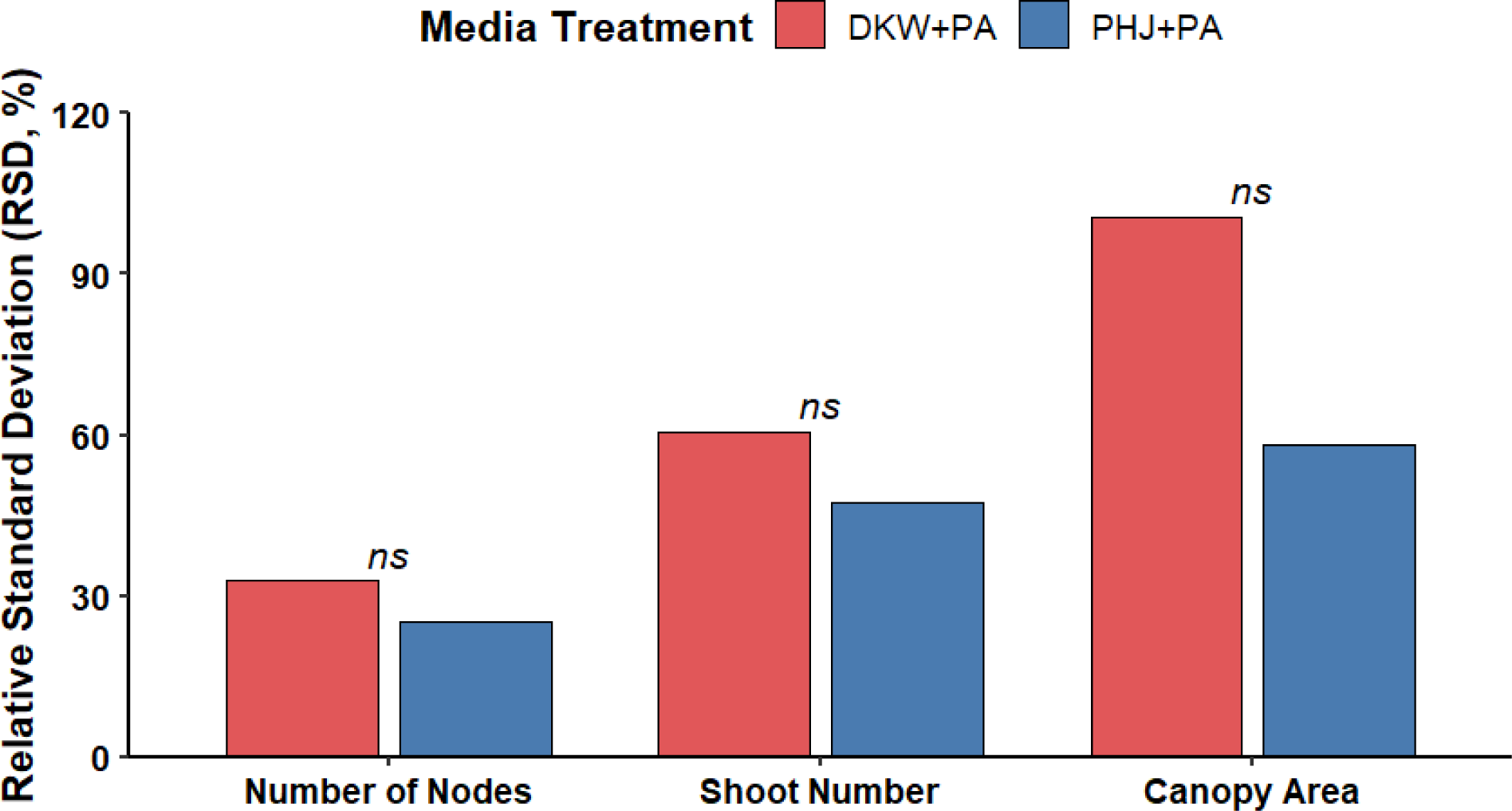
Comparison of overall plantlet uniformity following the subculture 1 phase. The RSD for DKW+PA and PHJ+PA are reported for each response variable quantified. Significance is noted with 95% confidence.

Plantlets responded better and more uniformly to PHJ+PA compared to DKW+PA. Plantlet sizes and morphologies were more uniform among ST and NC of each genotype when cultured on PHJ+PA. Swelling of the basal stem tissue leading to basal callus began to develop in the majority of replicates of each treatment, and the PhytoAx phenotype began to emerge in certain specimens. In general, PSS responded more appreciably to culture conditions than Roto (Fig. 13).

**Figure 13.**
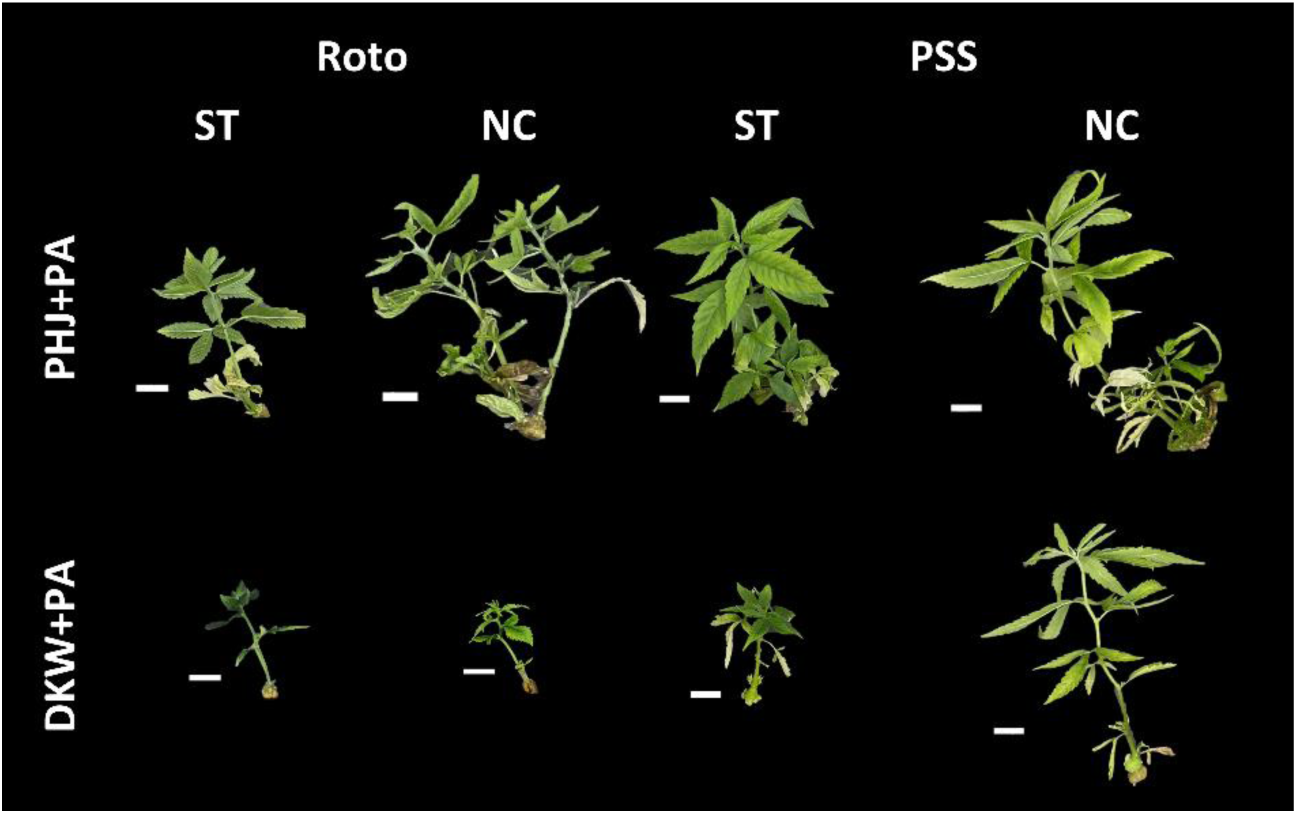
Representative specimens after the 4-week subculture 1 phase. Depicted are genotypes Roto and PSS grown from shoot tips (ST) and nodal cuttings (NC) on PHJ+PA and DKW+PA. Scale bars represent 10mm length.

#### 3.6.6. Subculture Phase 2 with PhytoAx

After 7-weeks in the subculture 2 phase, RSD trends continued to show greater uniformity of plantlets grown on PHJ+PA compared to DKW+PA for all the response variables tested. Number of nodes, shoot number, and canopy surface area of planets grown on PHJ+PA (21.31%, 47.25%, and 66.55%, respectively) were numerically more uniform than DKW+PA (28.46%, 62.61%, and 79.99%). However, the differences in relative uniformity remained insignificant (Fig. 14).

**Figure 14.**
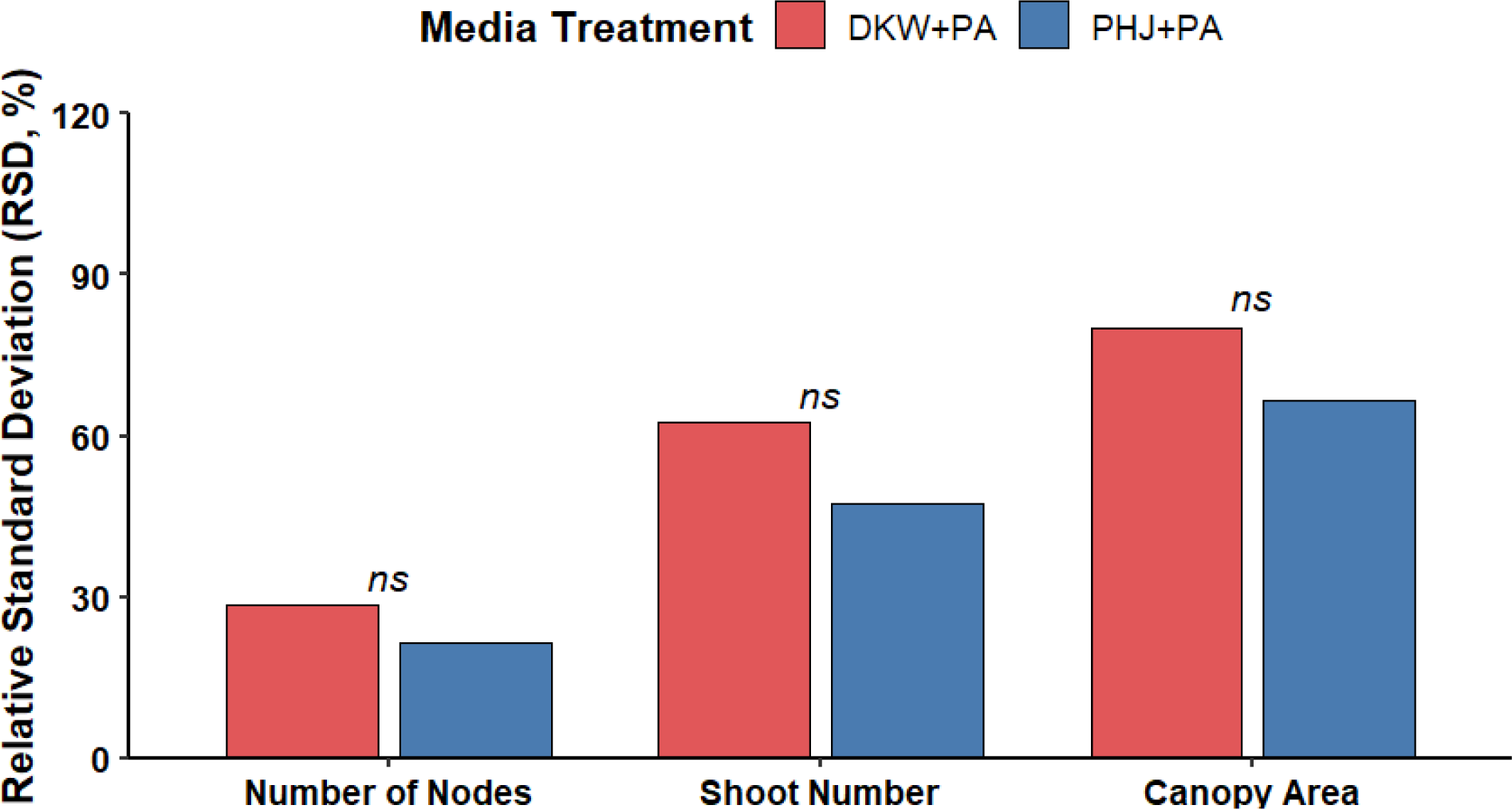
Comparison of overall plantlet uniformity following the subculture 2 phase. The RSD for DKW+PA and PHJ+PA are reported for each response variable quantified. Significance is noted with 95% confidence.

Plantlet response was more uniform on PHJ+PA compared to DKW+PA. Plantlet sizes and morphologies were more uniform among ST and NC of each genotype when cultured on PHJ+PA. In most cases, across genotypes and treatments, swelling of the basal stem resulted in the formation of basal callus. The PhytoAx phenotype was more dominant on DKW+PA compared to PHJ+PA and more pronounced in PSS compared to Roto. In general, PSS responded more appreciably to culture conditions than Roto (Fig. 15).

**Figure 15.**
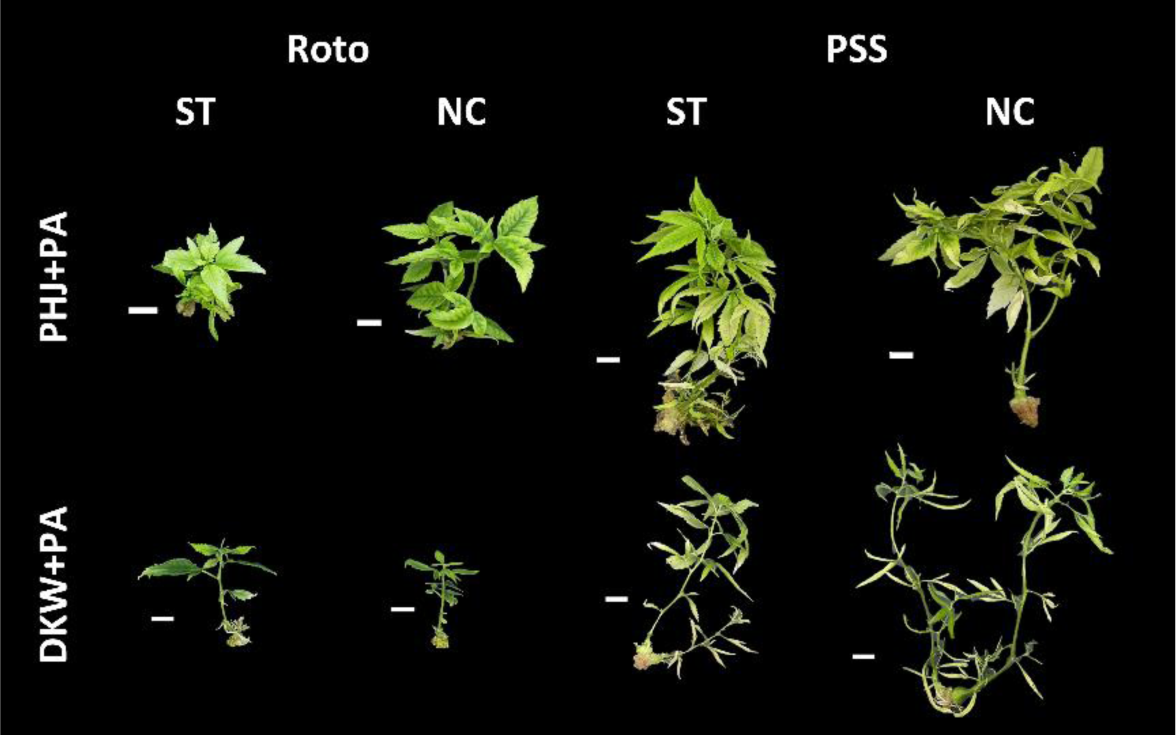
Representative specimens after the 7-week subculture 2 phase. Depicted are genotypes Roto and PSS grown from shoot tips (ST) and nodal cuttings (NC) on PHJ+PA and DKW+PA. Scale bars represent 10mm length.

#### 3.6.7. Subculture Phase 3 with PhytoAx

After 4-weeks in the subculture 3 phase, RSD showed that number of nodes of plantlets grown on PHJ+PA (13.75%) were significantly more uniform than plantlets grown on DKW+PA (30.29%). Shoot number, and canopy surface area of plantlets grown on PHJ+PA (37.32%, and 64.96%, respectively) were also more uniform than counterparts grown on DKW+PA (41.67%, and 77.10%, respectively. However, shoot number and canopy surface area were insignificant based on RSD (Fig. 16).

**Figure 16.**
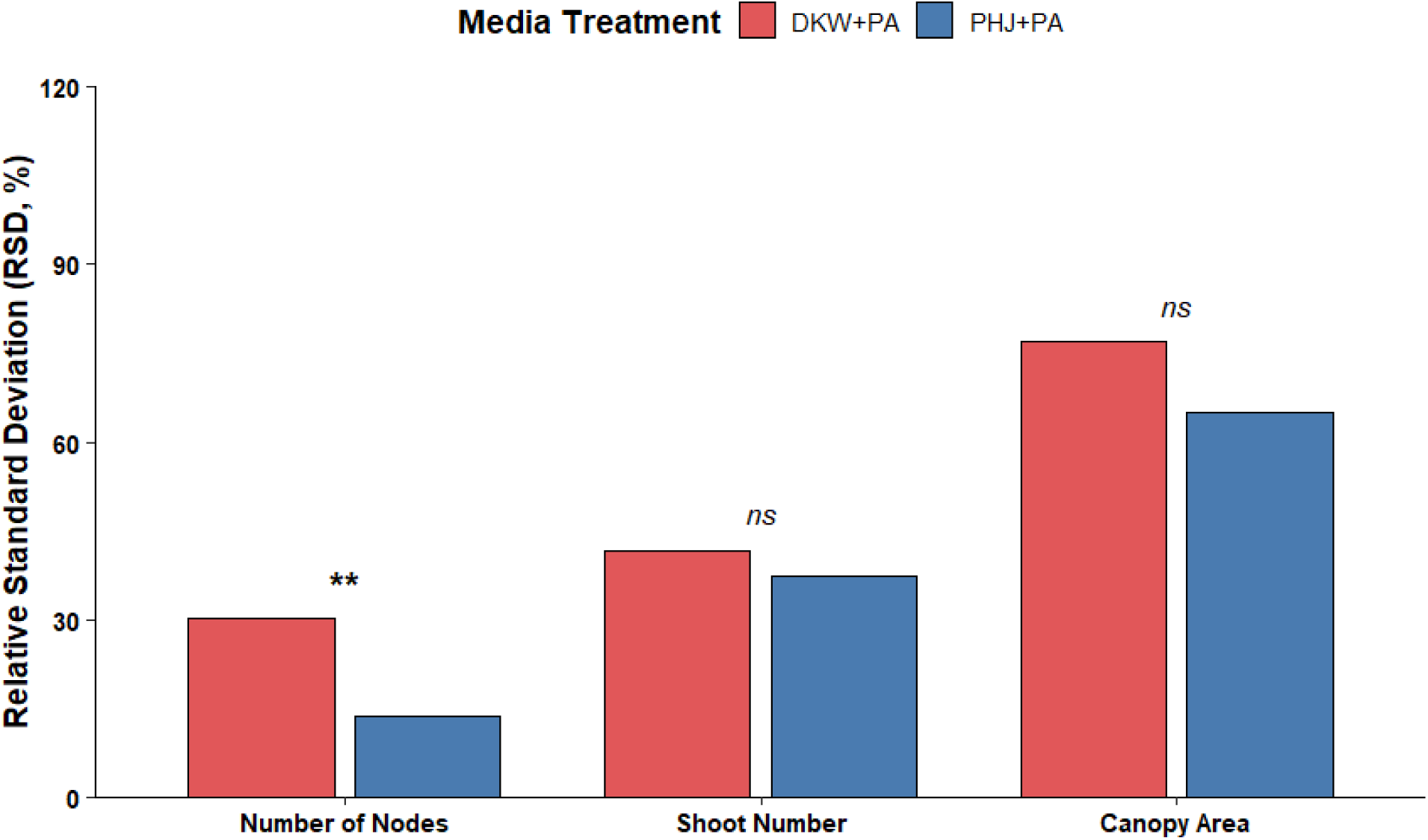
Comparison of overall plantlet uniformity following the subculture 3 phase. The RSD for DKW+PA and PHJ+PA are reported for each response variable quantified. Significance is noted with 95% confidence.

Plantlets were generally larger and more branched on PHJ+PA compared to DKW+PA. In most cases, across genotypes and treatments, plantlets appeared to revert from the PhytoAx phenotype to a more regular state, but retained basal callus and multiple shoot tips. This was generally more predominant on PHJ+PA, while plantlets on DKW+PA either retained their PhytoAx phenotype or appeared unresponsive to PhytoAx at this stage. In general, PSS responded more appreciably to culture conditions than Roto (Fig. 17).

**Figure 17.**
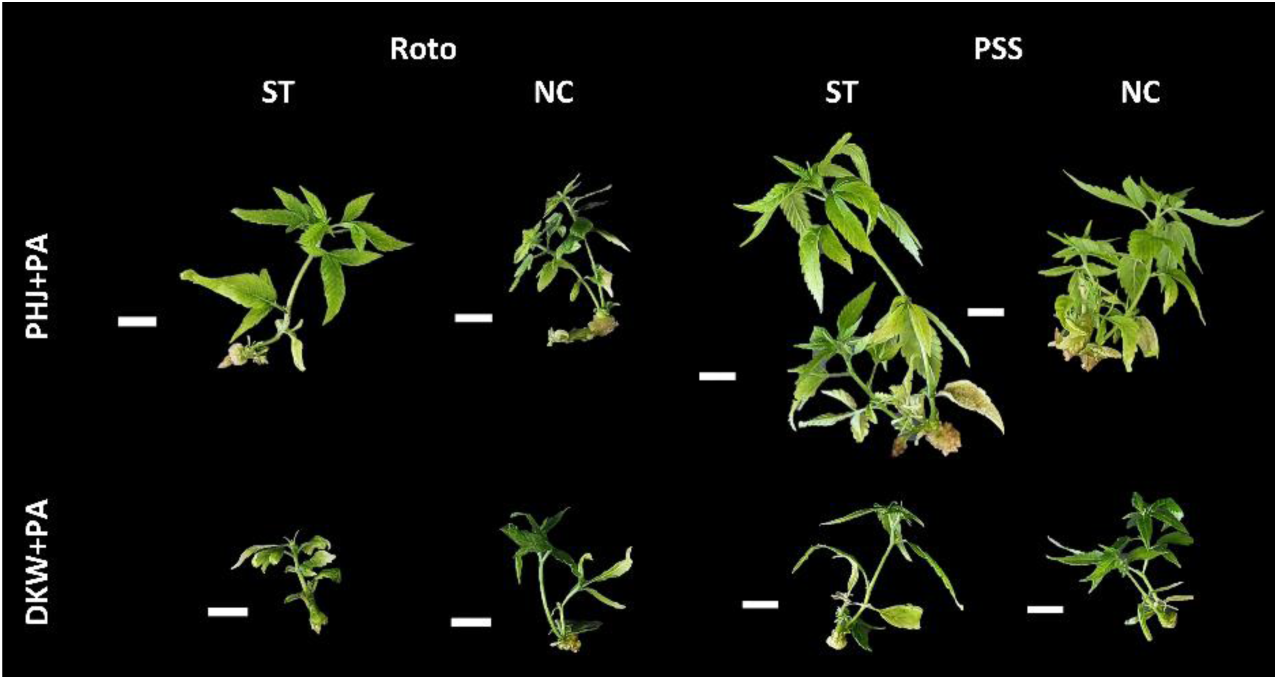
Representative specimens after the 4-week subculture 3 phase. Depicted are genotypes Roto and PSS grown from shoot tips (ST) and nodal cuttings (NC) on PHJ+PA and DKW+PA. Scale bars represent 10mm length.

#### 3.6.8. Effect of Media with PhytoAx Throughout Subculture

Following the first round of subculture, PHJ+PA NC, produced significantly greater shoot numbers and number of nodes compared to explants on basal PHJ (S4). Shoot numbers were significantly greater on DKW+PA for both explant types compared to basal DKW (S5). No other significant differences related to the presence of PhytoAx occurred from the first round of subculture.

The second round of subculture resulted in significantly greater shoot numbers for both explant types on PHJ+PA compared to basal PHJ. Number of nodes was significantly greater for PHJ+PA NC. Canopy surface area was significantly greater for PHJ+PA ST (S6). Significantly greater shoot numbers were produced on DKW+PA compared to basal DKW for both explant types (S7). No other significant differences related to the presence of PhytoAx occurred from the second round of subculture.

The third round of subculture allowed significantly greater shoot numbers on DKW+PA for both explant types compared to basal PHJ. Significantly greater number of nodes occurred on PHJ+PA NC. Canopy surface area was significantly greater on PHJ+PA for NC (S8). Number of nodes was significantly greater for basal DKW ST (S9). No other significant differences related to the presence of PhytoAx occurred from the third round of subculture.

### 3.7. Culture Initiation Experiment

Shoot numbers and number of nodes were significantly greater for SLH on PHJ compared to DKW on the third round of subculture following culture initiation (S10). No significant differences in SLH shoot numbers or node numbers occurred when cultured on PHJ+PA or DKW+PA following three rounds of subculture after culture initiation. Number of nodes was significantly greater for TRS10 cultures on PHJ+PA compared to DKW+PA. No significant differences in shoot numbers occurred for TRS10 (S11).

## 4. Discussion

The current investigation represents one of the most comprehensive media optimization studies in recent history. The resulting PHJ media was specifically developed to accommodate a wide array of cannabis cultivars to address the well documented challenges associated with *in vitro* growth and genotypic variability throughout cannabis micropropagation. Many media optimization studies focus on the separate optimization of individual media components or grouped categories of components (i.e. macronutrients, mesonutrients, and micronutrients) (Bao et al., 2025; Nor et al., 2017; Zarei et al., 2023), which can often not capture the mixed effects of multiple factors on plant responses (Pepe et al., 2025b). The current study focused on the concurrent optimization of individual media components to help overcome the setbacks of the aforementioned approaches. The resulting PHJ media represents a unique formulation when compared to the existing alternatives, which was made possible with the use of ML to help recognize complex interaction patterns and their effects on cannabis growth and development. Specifically, three machine learning algorithms (GRNN, SVR, and ANFIS) and a bagging-based ensemble model were used to capture the relationships among media components and key morpho-physiological traits of cannabis explants. Plant growth responses are governed by tightly coordinated physiological processes, including ion uptake, osmotic regulation, enzyme activation, and metabolic fluxes, all of which are influenced by both the concentration and balance of macro- and micro-nutrients (Pasternak and Steinmacher, 2024). These interdependencies create a multidimensional response surface that is difficult to describe using linear or single-model approaches (Arab et al., 2018; Sadat-Hosseini et al., 2023). The complex and nonlinear nature of plant-nutrient integrations occurring *in vitro* can predictably be captured using ML approaches (Arteta et al., 2022; Hameg et al., 2020), and in the current case was successfully captured by the ability of the developed ensemble model.

While all tested models were able to establish predictive relationships to varying degrees, the ensemble approach, which combined the various models, provided improved predictive performance compared to any of the individual models. This indicates that the mineral nutrients collectively contributed to shoot length, node number, and canopy surface area variability. The integration of the diverse learning patterns more accurately captured the complex dynamics for superior performance (Ngo et al., 2022; Wu et al., 2018). While GRNN is particularly effective in modeling smooth nonlinear relationships due to its probabilistic nature, SVR is designed to maximize generalization by balancing model complexity and prediction error (Hesami and Jones, 2020). Alternatively, ANFIS incorporates rule-based fuzzy logic, allowing human-like reasoning to approximate systems with uncertainty (Ghosh et al., 2026). However, each of these models is constrained by its underlying assumptions and sensitivity to data structure (Barbierato and Gatti, 2024). By aggregating predictions through bagging, the ensemble approach can reduce variance and mitigate individual model biases, leading to a more stable and generalized representation of the underlying biological system (Hesami et al., 2020). This is especially relevant in plant tissue culture studies, where experimental noise and biological variability are often unavoidable (Narra et al., 2025; Zhou et al., 2023).

Consistent with a growing body of evidence (Hesami and Jones, 2020; Narra et al., 2025), the current findings reinforce the utility of ML-based optimization for plant tissue culture. Traditional statistical approaches are often inadequate for resolving high-order interactions among factors, especially when concurrently optimizing numerous nutrients, whereas data-driven models have demonstrated improved flexibility and predictive accuracy (Arab et al., 2018; Sadat-Hosseini et al., 2023). Similar improvements using ensemble techniques that combine multiple algorithms to enhance robustness and reduce overfitting have been used for pomegranate proliferation (Zarbakhsh et al., 2024) and chrysanthemum somatic embryogenesis (Hesami et al., 2020). However, some studies report only marginal gains using ensemble methods, suggesting that their effectiveness may depend on dataset size, feature diversity, and the degree of nonlinearity in the system (Fan et al., 2025; Shaikh et al., 2024; Tong and Li, 2025). In this context, the results presented strengthen the value of ensemble learning while highlighting the importance of selecting complementary base models (Zhou et al., 2023).

The Pareto-optimal solution, PHJ, was derived from the NSGA-II multi-objective optimization and represents a targeted strategy for maximizing key cannabis growth and development parameters by selectively enriching both macro- and micro-nutrients to support crucial physiological processes. This exemplifies how evolutionary algorithms can identify precise macro- and micro-nutrient combinations that leverage specific mechanistic pathways to enhance micropropagation efficiency for cannabis (Hesami and Jones, 2020; Narra et al., 2025). Despite these advantages, it is important to note that ML models remain empirical tools that do not inherently reveal causal mechanisms (Pepe et al., 2025b; Zhou et al., 2023). While they can approximate complex relationships with high accuracy, their predictions must be interpreted within the framework of plant physiology and validated experimentally Pepe et al., 2025b).

The media resulting from the aforementioned ML approach is unique in many ways when compared to existing alternatives. For instance, PHJ contains 7.96 g/L total nutrient salts. This is 53% more than the total nutrient salts in DKW, and 84% more than MS. Cannabis has high metabolic activity (Livingston et al., 2022; Pepe et al., 2022), which can benefit from increased nutrient availability *in vitro* (Lubell-brand et al., 2021; Zarei et al., 2023). The high concentrations of nutrient salts available in the optimized PHJ medium is a direct reflection of this. While PHJ contains 46% more NH_4_NO_3_ than DKW, and 25% more than MS, which is among the most important macronutrient for plants in culture, perhaps the most striking modifications are seen with the relative differences of micronutrients. For instance, relative amounts of FeSO_4_, CuSO_4_, MnSO_4_, Na_2_MoO_4_, Zn(NO_3_)_2_, NiSO_4_, and H_3_BO_3_ in PHJ are 49%, 338%, 400%, 368%, 412%, 480%, and 4.48% higher than in DKW, respectively. Similarly, relative amounts of FeSO_4_, CuSO_4_, MnSO_4_, and Zn(NO_3_)_2_ in PHJ are 81%, 42.82%, 892%, and 911% higher than in MS, respectively. With this in perspective, the ML approach has afforded an extremely unique culture media for cannabis what would likely not have been developed using traditional approaches.

Another powerful tool afforded by ML is the sensitivity analysis, a type of feature importance analysis that evaluates the degree with which a model’s output varies in response to alterations to the input features. In reference to the current work, the feature importance evaluates the relative impact each salt has on the specific growth parameters. Specifically, the relative influence of nutrients was determined by calculating VSR to evaluate their importance on each trait. Interestingly, VSR suggests that many of the most influential media components for shoot length, number of nodes, and canopy surface area were chelating agents and/or micronutrients. For example, the top three for shoot length were Na_2_EDTA, Zn(NO_3_)_2_, and Na_2_MoO_4_, for number of nodes were KSO_4_, Na_2_MoO_4_, and MnSO_4_, and for canopy surface area were CaCl_2_, Zn(NO_3_)_2,_ and H_3_BO_3_. Chelating agents such as EDTA help maintain the solubility and bioavailability of certain micronutrients, such as iron, which along with zinc and manganese, are essential for chlorophyll synthesis, photosystem performance, oxidative stress management, and enzyme regulation (Arinaitwe et al., 2026; Jomova et al., 2025; Madhupriyaa et al., 2024; Vyskot and Bezdek, 1984). Molybdenum function as a cofactor for nitrogen metabolism (Kamal et al., 2023; Saleem et al., 2020). Thus, Na_2_EDTA would function to improve bioavailability of these micronutrient metals (Kamal et al., 2023; Saleem et al., 2020) that serve as enzymatic control points for regulatory roles in plant growth responses (Bhatla and Kathpalia, 2023). The findings surrounding the relative importance of micronutrients and chelating agents exemplifies that while micronutrients are only required in small amounts, their physiological functions are critical to the growth and wellbeing of plants. Further, this highlights the deficiency in media optimization approaches where micronutrients are either overlooked or treated as a single group with static ratios (Bao et al., 2025; Hameg et al., 2020; Zarei et al., 2023). As a result, many of the existing media available predominantly differ in macronutrient levels but are very similar in respect to micronutrient composition, both quantitatively and qualitatively. Based on the current results, micronutrient optimization is likely more important for future media optimization studies than is generally acknowledged.

The comparatively lower VSR of certain media components suggests that not all nutrients exert equal control over the modeled traits within the tested concentration ranges, but this should not be interpreted as a lack of importance. Rather, this likely reflects either sufficient baseline availability, redundancy in physiological function, or limited involvement in the specific developmental pathways measured (Jamshidi et al., 2016). For example, if a larger range of nutrient concentrations that fall outside of the optimal window were tested, VSR would identify their relative importance as much greater. As such, sensitivity analysis should be interpreted carefully and can simply indicate that these components are least optimized, rather than the most important from a physiological standpoint, and should potentially be the focus of future studies. This also highlights a key limitation of nutrient optimization studies, as the effect of a given element cannot be fully understood in isolation, since synergistic and antagonistic interactions often modulate overall plant responses (Pepe et al., 2025b). These results align with the broader understanding that micronutrients and their availability often have disproportionately large effects relative to their concentrations (Arab et al., 2018; Jamshidi et al., 2016; Sadat-Hosseini et al., 2023). The degree of sensitivity presented includes both macro- and micro-nutrients, highlighting that all essential elements are critical and highly interconnected, resulting in multiple elements simultaneously influencing growth (Hesami et al., 2020).

It has been reported that DKW offers an improved nutrient profile for cannabis relative to MS (Kastelec et al., 2025; Page et al., 2021), likely due to higher concentration of macro- and micro-elements, including metals (Kastelec et al., 2025). While this was not observed in validation experiment 2, where MS largely performed better than DKW, a key distinction is that Page et al., (2021) conducted their study with the addition of TDZ in the media, resulting in extreme callusing, hyperhydricity, and culture decline. When they cultured plants without TDZ, the side effects were less pronounced (Page et al., 2021), suggesting that the challenge with MS basal media involved an interaction between the media composition and the PGR rather than being a straight forward issue related exclusively to plant nutrition. This, along with a high degree of genotypic variability, contributes to some apparent contradictions in the literature in which certain media containing MS outperform DKW (Holmes et al., 2021). While these inconsistent results can complicate interpretations, when evaluated in context, they appear consistent with our general understanding of the interactions of media components and impacts on plant responses.

Interestingly, DKW was originally developed for walnut micropropagation to address similar challenges to those observed in cannabis, including long term culture decline (Driver and Kuniyuki, 1984). By adjusting various media components, Driver and Kuniyuki (1984) were able to overcome these issues and establish an effective micropropagation system for *Juglans hindsii* x *J. regia*. For cannabis, the implementation of DKW overcame many associated challenges to facilitate successful micropropagation, although certain nutrient imbalances remained (Page et al., 2021). While the nutrient profile afforded by DKW can improve cannabis micropropagation in some instances, the current results show that DKW remains suboptimal for cannabis and cannabis growth can be enhanced by increasing the relative amounts of certain media components to improve nutrient ratios for a highly metabolic plant. This targeted nutrient enrichment likely ensures that the metabolic machinery required for both node formation and canopy expansion operate at optimal capacity. As DKW media was developed for walnut, its broad suitability for cannabis forecasts the further suitability of PHJ to be used on other plant species with similar challenges and nutritional requirements.

Cannabis can be notoriously recalcitrant *in vitro* due to high genotypic variation that prevents the implementation of a universal culture protocol (Kastelec et al., 2025; Monthony et al., 2021; Page et al., 2021). This is demonstrated in the current study, where cultivar response significantly differs on a particular media (for instance on MS), indicating that the best media often depends on the cultivar in question. Universal protocols are rarely guaranteed in the field of plant tissue culture, since genetic, epigenetic, and physiological factors contribute to variable plantlet performance, often even among specimens of the same cultivar (Hesami et al., 2025). However, the combined data from the current investigation validates the use of PHJ to reliably allow appreciable, uniform growth and developmental responses for the nine cannabis cultivars used for its development and testing. Of particular note, PHJ not only resulted in better average growth across genotypes, but the inter-cultivar variability was generally lower. This is noticeable when examining RSD and growth results in validation experiment 2 and in the subculture experiment both with and without PhytoAx. While certain cultivars exhibited better responses to culture in general, the overarching trends in the current work demonstrates that PHJ allows superior performance compared to both DKW and MS. This trend remains consistent for culture initiation and over multiple rounds of subculture.

During the subculture experiment, although the initial introduction of explants to PHJ allowed greater number of nodes, number of shoots, and canopy surface areas, it did not yield immediate uniformity based on genotype and explant type. However, the progressive reduction in RSD of PHJ compared to DKW was more pronounced in subculture phase 2, and continued to progress though to subculture phase 3, especially for number of nodes and canopy surface area. For number of shoots, uniformity on PHJ began relatively low, then increased after subculture phase 2, and remained fairly stable though subculture phase 3. This delayed response is attributed to a physiological carryover effect of plantlets being maintained on DKW prior to their transfer to PHJ for experimentation. Plantlets likely require multiple culture passes to acclimate to the new nutritional profile of PHJ before more stable uniformity can be achieved (Al-Oqab et al., 2023; Hesami et al., 2023). Number of nodes reflects meristem activity and developmental patterning processes that may be less immediately responsive to short-term nutrient fluctuations (Ristova et al., 2016). Canopy surface area integrates both structural and spatial growth components, making it particularly sensitive to cumulative effects of nutrient interactions over time (Burgess et al., 2021). Higher concentrations of calcium and potassium available in PHJ may have contributed to the initial variability, ultimately leading to higher uniformity. Calcium is integral to cell wall stabilization and acts as a secondary messenger in signal transduction pathways (Pepe et al., 2025b), which may explain its strong association with traits involving structural expansion, such as canopy surface area. Potassium regulates osmotic balance and stomatal function, influencing turgor-driven growth processes, while phosphorus is central to energy transfer and nucleic acid synthesis (Pasternak and Steinmacher, 2024). Ultimately, the trends demonstrate that PHJ allows immediate overall improvement in response variables when transferred from an alternative basal media. However, uniformity among cultivars and explant types requires multiple rounds of sequential subcultures following the initial transfer. This highlights a trait-specific timeline for full tissue acclimation.

The addition of PhytoAx to PHJ resulted in significantly greater shoot numbers for both tested explant types and significantly greater number of nodes and canopy surface areas for NC compared to PHJ alone at the end of subculture phase 3. Alternatively, the addition of PhytoAx to DKW did not significantly increase any of the measured plantlet responses for either explant type by the end of subculture phase 3. Additionally, phenotypic uniformity was consistently superior on PHJ+PA compared to DKW+PA. The immediate and consistent uniformity among explant type and genotype largely stems from a synergistic interaction between the PHJ nutrient profile and the added PGRs. Although the addition of exogenous PGRs can often induce or exacerbate the occurrence of morpho-physiological disorders (Barua et al., 2025; Zunazri et al., 2024), the optimized nutrient composition of PHJ effectively mitigated these occurrence and perpetuation of the PhytoAx Phenotype. As a result, the nutrient-PGR compatibility afforded by PHJ+PA allowed the plantlets to more effectively use PhytoAx, facilitating uniform growth responses across explant types and cultivars, beginning with subculture phase 1. For number of nodes and shoot number, this uniformity persisted and progressively improved over subculture phase 2, reaching its peak by the end of subculture phase 3. Interestingly, by the end of subculture phase 3, RSD values between PHJ and PHJ+PA for both number of nodes (13.80% and 13.75%, respectively) and shoot number (37.45% and 37.32%) were nearly identical. Ultimately, this convergence exemplifies that sequential subculturing on PHJ and PHJ+PA enables tissues to achieve a uniform physiological equilibrium, proving that prolonged phenotypic stability is achievable with optimization of basal nutrients, regardless of the presence of PGRs.

While there are contradictory reports, it has been posited that MS is not ideal for *in vitro* cannabis growth (Page et al., 2021; Zarei et al., 2023) and an improved DKW-based medium would better facilitate growth for micropropagation of cannabis (Page et al., 2021). As noted above, the high concentrations of specific macro- and micro-nutrients afforded by DKW enables more appreciable growth responses for micropropagated cannabis compared to other commonly used basal formulations such as MS (Kastelec et al., 2025). The optimized PHJ media takes this one step further and finetunes nutrient concentrations for cross-cultivar optimization of multiple growth parameters, demonstrating how concurrent nutrient optimization for multi-objective response variables can simultaneously tailer nutrient balances to improve overall plant quality. Potential for a reduction in the number of nodes produced over successive subcultures has been associated with cannabis cultured on DKW (Kastelec et al., 2025), which was observed in the current study. When PHJ was tested for longer term performance over multiple subcultures, the decrease in node count was less pronounced compared to DKW. Furthermore, PHJ+PA mitigated the reduction of node number, whereas adding DKW+PA amplified the reduction of node number on subculture 3. This highlights the importance of optimizing basal salt formulations before plant PGRs. Possible causes of the observed decline include the accumulation of epigenetic or physiological stress (Kastelec et al., 2025), which may be alleviated with the appropriate concentration and ratios of mineral salts (Hameg et al., 2020; Pasternak and Steinmacher, 2024; Pepe et al., 2025a). Additionally, PHJ contains CoCl_2_ and KI, which are absent from DKW. While PHJ+PA significantly improved shoot multiplication compared to DKW+PHJ over subcultures 2 and 3 it is recommended that explants be moved to PHJ with the absence of PA for rooting and acclimation due to possible perpetuation of the PhytoAx phenotype.

For the subculture experiment, PHJ allowed significantly greater shoot number and number of nodes compared to DKW, whereas only number of nodes was significantly greater with PHJ+PA compared to DKW+PA for one cultivar tested. However, in all cases, PHJ -based media outperformed DKW -based media, and media with PhytoAx outperformed those without. The collection, sectioning, and sterilization of explants for culture initiation causes severe tissue stress, while the addition of hormones and high concentrations of nutrients can further perpetuate stress signaling and force cell division before tissue acclimation (Liu et al., 2024). This drastically reduces the overall performance of explants and causing morpho-physiological disorders (Martini et al., 2022). While the current study scratches the surface on showing the benefit of PHJ with and without PGRs for culture initiation, follow-up work must be conducted to further confirm the efficacy of PHJ at different concentrations, with and without PGRs for culture initiation.

Optimizing the synergy between plant PGRs and mineral salt solutions has long been a focal point of *in vitro* cannabis research. These investigations often prioritize the refinement of both mineral salts and phytohormones simultaneously (Baek et al., 2024; Das et al., 2024), or focus specifically on isolating the most effective concentrations of PGRs alone (Johnson et al., 2025; Kastelec et al., 2025). The success of the micropropagation puzzle is multifactorial, nonlinear, and is difficult to model using traditional techniques, since mineral nutrients show dynamic interactions that impact culture success (Pepe et al., 2025a). Moreover, phytohormone efficacy is largely interactional based on the concentrations and ratios of the additives (Thacker et al., 2018), and their further interactions with media nutrient profiles (Page et al., 2021). The development of PHJ focused specifically on the optimization of mineral salts to enhance *in vitro* performance for cannabis micropropagation. Benefits were further compounded when PHJ was paired with PhytoAx, a previously optimized, proprietary phytohormone solution. A modular approach involving the independent optimization of nutrient components and phytohormones, followed by integration and subsequent synergistic refinement offers a promising framework for future micropropagation work across diverse species. The efficacy of PHJ+PA highlights the potential of PHJ in diverse plant tissue culture applications beyond microproagation, including meristem culture, callogenesis, or somatic embryogenesis. Future work should prioritize integrating PHJ with established protocols to fully assess its broad scale applicability.

While it is commonly accepted that basal media formulations should be specifically engineered to suit the genotype of interest (Espinosa-Leal et al., 2018; Greenway et al., 2012), there exist various media with broad applicability for numerous plants (Greenway et al., 2012; Kastelec et al., 2025; Page et al., 2021). Furthermore, while some plants benefit from lower overall concentrations of mineral salts *in vitro* (Fadel et al., 2010; Kumar et al., 2024; Tinoammini et al., 2024), other plants thrive from the availability of enhanced concentrations (Jain et al., 2012; Lekamge et al., 2021; Wada et al., 2013). Given the efficacy of PHJ on a wide array of cannabis cultivars, there is high probability for its broad utility with a variety of plant species. Future research focusing on the performance of PHJ for *in vitro* applications across diverse taxa is needed, and the incorporation of different media strengths and PGR concentrations is recommended. This approach mirrors the use of highly successful basal media formulations like MS and DKW, which have achieved widespread versatility across a variety of plants.

## 5. Conclusion

The ML-mediated development of PHJ represents one of the most comprehensive media optimization endeavours thus far. Use of a hybridized ensemble-NSGA-II approach resulted in the development of a completely unique basal media formulation with exceptional ratios of mineral salts. This approach outperforms methods traditionally used for plant tissue culture optimization by identifying complex, collective nutrient interactions, with ensemble approaches to allow superior predictive performance for concurrent optimization of individual media components. The resulting PHJ media allows improved and uniform plantlet responses for key micropropagation parameters, including number of nodes, shoot number, and canopy surface area for the nine cultivars used in its development and validation. These results remained consistent for multiple micropropagation stages, from explant initiation though multiple subculture phases. Furthermore, robust performance of PHJ, both with and without PGRs, underscores its potential versatility for diverse plant tissue culture applications beyond standard micropropagation. The ability of PHJ to successfully overcome genotypic recalcitrance, improve growth parameters, and maintain uniformity across multiple cultivars is telling of its potential applicability with an assortment of plant species beyond cannabis.

## Author Contributions

The work was conceptualized by Marco Pepe, Mohsen Hesami, and A.M.P. Jones. Experiments were conducted by Marco Pepe and Mohsen Hesami. Data was processed, analyzed, and visualized by Marco Pepe and Mohsen Hesami. The project was supervised by A.M.P. Jones. The original manuscript draft was drafted by Marco Pepe and Mohsen Hesami. The manuscript was reviewed by A.M.P. Jones and edited by Marco Pepe. All authors have read the current manuscript and have agreed upon publication in its current form.

## Acknowledgments

The authors want to acknowledge and thank Evan Kay for his support with technical aspects of the work conducted.

## Data Availability Statement

The current manuscript reports only original data. All data is included in the article and/or supplementary materials. Further inquiries can be directed to the corresponding author.

