## Supplementary material for "Introducing PHJ Media: A Unique Machine Learning -Driven Basal Formulation to Overcome Recalcitrance for Multi-Genotype Micropropagation of *Cannabis sativa* L": S1, S2, S3, S4, S5, S6, S7, S8, S9, S10, S11

S1. Salt components included in treatments for the initial experiment. All values represent mg/L. Additional components of these treatments included 100mg/L Myo-Inositol, 2mg/L Glycine, 1mg/L Nicotinic acid, 2mg/L Thiamine, 0.004mg/L CoCl2, and 0.1mg/L KI, which were added consistently to all treatments. Additionally, each treatment included 0.6% (w/v) agar, 3% (w/v) sucrose, and pH = 5.7.

| Treatment | NH4NO3 | K2SO4 | Ca(NO3)2 | CaCl2 | KH2PO4 | MgSO4 | FeSO4 · 7H2O | Na2EDTA · 2H2O | CuSO4 · 5H2O | MnSO4 · H2O | Na2MoO4 · 2H2O | Zn(NO3)2.6H2O | NiSO4.6H2O | H3BO3 |
|---|---|---|---|---|---|---|---|---|---|---|---|---|---|---|
| DKW | 1416 | 1559 | 1367 | 112.5 | 265 | 361.49 | 33.8 | 45.4 | 0.25 | 33.5 | 0.39 | 17 | 0.005 | 4.8 |
| 1 | 708 | 1559 | 1367 | 112.5 | 265 | 361.49 | 33.8 | 45.4 | 0.25 | 33.5 | 0.39 | 17 | 0.005 | 4.8 |
| 2 | 708 | 779.5 | 683.5 | 112.5 | 265 | 361.49 | 33.8 | 45.4 | 0.25 | 33.5 | 0.39 | 17 | 0.005 | 4.8 |
| 3 | 708 | 2338.5 | 2050.5 | 112.5 | 265 | 361.49 | 33.8 | 45.4 | 0.25 | 33.5 | 0.39 | 17 | 0.005 | 4.8 |
| 4 | 708 | 1559 | 1367 | 56.25 | 132.5 | 180.745 | 33.8 | 45.4 | 0.25 | 33.5 | 0.39 | 17 | 0.005 | 4.8 |
| 5 | 708 | 1559 | 1367 | 168.75 | 397.5 | 542.235 | 33.8 | 45.4 | 0.25 | 33.5 | 0.39 | 17 | 0.005 | 4.8 |
| 6 | 708 | 1559 | 1367 | 112.5 | 265 | 361.49 | 16.9 | 22.7 | 0.25 | 33.5 | 0.39 | 17 | 0.005 | 4.8 |
| 7 | 708 | 1559 | 1367 | 112.5 | 265 | 361.49 | 50.7 | 68.1 | 0.25 | 33.5 | 0.39 | 17 | 0.005 | 4.8 |
| 8 | 708 | 1559 | 1367 | 112.5 | 265 | 361.49 | 33.8 | 45.4 | 0.125 | 16.75 | 0.195 | 8.5 | 0.0025 | 2.4 |
| 9 | 708 | 1559 | 1367 | 112.5 | 265 | 361.49 | 33.8 | 45.4 | 1.25 | 167.5 | 1.95 | 85 | 0.025 | 24 |
| 10 | 708 | 1559 | 1367 | 112.5 | 265 | 361.49 | 33.8 | 45.4 | 5 | 670 | 7.8 | 340 | 0.1 | 96 |
| 11 | 708 | 1559 | 1367 | 112.5 | 265 | 361.49 | 33.8 | 45.4 | 5 | 670 | 7.8 | 340 | 0.1 | 480 |
| 12 | 2124 | 1559 | 1367 | 112.5 | 265 | 361.49 | 33.8 | 45.4 | 0.25 | 33.5 | 0.39 | 17 | 0.005 | 4.8 |
| 13 | 2124 | 779.5 | 683.5 | 112.5 | 265 | 361.49 | 33.8 | 45.4 | 0.25 | 33.5 | 0.39 | 17 | 0.005 | 4.8 |
| 14 | 2124 | 2338.5 | 2050.5 | 112.5 | 265 | 361.49 | 33.8 | 45.4 | 0.25 | 33.5 | 0.39 | 17 | 0.005 | 4.8 |
| 15 | 2124 | 1559 | 1367 | 56.25 | 132.5 | 180.745 | 33.8 | 45.4 | 0.25 | 33.5 | 0.39 | 17 | 0.005 | 4.8 |
| 16 | 2124 | 1559 | 1367 | 168.75 | 397.5 | 542.235 | 33.8 | 45.4 | 0.25 | 33.5 | 0.39 | 17 | 0.005 | 4.8 |
| 17 | 2124 | 1559 | 1367 | 112.5 | 265 | 361.49 | 16.9 | 22.7 | 0.25 | 33.5 | 0.39 | 17 | 0.005 | 4.8 |
| 18 | 2124 | 1559 | 1367 | 112.5 | 265 | 361.49 | 50.7 | 68.1 | 0.25 | 33.5 | 0.39 | 17 | 0.005 | 4.8 |
| 19 | 2124 | 1559 | 1367 | 112.5 | 265 | 361.49 | 33.8 | 45.4 | 0.125 | 16.75 | 0.195 | 8.5 | 0.0025 | 2.4 |
| 20 | 2124 | 1559 | 1367 | 112.5 | 265 | 361.49 | 33.8 | 45.4 | 1.25 | 167.5 | 1.95 | 85 | 0.025 | 24 |
| 21 | 2124 | 1559 | 1367 | 112.5 | 265 | 361.49 | 33.8 | 45.4 | 5 | 670 | 7.8 | 340 | 0.1 | 96 |
| 22 | 2124 | 1559 | 1367 | 112.5 | 265 | 361.49 | 33.8 | 45.4 | 5 | 670 | 7.8 | 340 | 0.1 | 480 |
| 23 | 1416 | 779.5 | 683.5 | 112.5 | 265 | 361.49 | 33.8 | 45.4 | 0.25 | 33.5 | 0.39 | 17 | 0.005 | 4.8 |
| 24 | 1416 | 779.5 | 683.5 | 56.25 | 132.5 | 180.745 | 33.8 | 45.4 | 0.25 | 33.5 | 0.39 | 17 | 0.005 | 4.8 |
| 25 | 1416 | 779.5 | 683.5 | 168.75 | 397.5 | 542.235 | 33.8 | 45.4 | 0.25 | 33.5 | 0.39 | 17 | 0.005 | 4.8 |
| 26 | 1416 | 779.5 | 683.5 | 112.5 | 265 | 361.49 | 16.9 | 22.7 | 0.25 | 33.5 | 0.39 | 17 | 0.005 | 4.8 |
| 27 | 1416 | 779.5 | 683.5 | 112.5 | 265 | 361.49 | 50.7 | 68.1 | 0.25 | 33.5 | 0.39 | 17 | 0.005 | 4.8 |
| 28 | 1416 | 779.5 | 683.5 | 112.5 | 265 | 361.49 | 33.8 | 45.4 | 0.125 | 16.75 | 0.195 | 8.5 | 0.0025 | 2.4 |
| 29 | 1416 | 779.5 | 683.5 | 112.5 | 265 | 361.49 | 33.8 | 45.4 | 1.25 | 167.5 | 1.95 | 85 | 0.025 | 24 |
| 30 | 1416 | 779.5 | 683.5 | 112.5 | 265 | 361.49 | 33.8 | 45.4 | 5 | 670 | 7.8 | 340 | 0.1 | 96 |
| 31 | 1416 | 779.5 | 683.5 | 112.5 | 265 | 361.49 | 33.8 | 45.4 | 5 | 670 | 7.8 | 340 | 0.1 | 480 |
| 32 | 1416 | 2338.5 | 2050.5 | 112.5 | 265 | 361.49 | 33.8 | 45.4 | 0.25 | 33.5 | 0.39 | 17 | 0.005 | 4.8 |
| 33 | 1416 | 2338.5 | 2050.5 | 56.25 | 132.5 | 180.745 | 33.8 | 45.4 | 0.25 | 33.5 | 0.39 | 17 | 0.005 | 4.8 |
| 34 | 1416 | 2338.5 | 2050.5 | 168.75 | 397.5 | 542.235 | 33.8 | 45.4 | 0.25 | 33.5 | 0.39 | 17 | 0.005 | 4.8 |
| 35 | 1416 | 2338.5 | 2050.5 | 112.5 | 265 | 361.49 | 16.9 | 22.7 | 0.25 | 33.5 | 0.39 | 17 | 0.005 | 4.8 |
| 36 | 1416 | 2338.5 | 2050.5 | 112.5 | 265 | 361.49 | 50.7 | 68.1 | 0.25 | 33.5 | 0.39 | 17 | 0.005 | 4.8 |
| 37 | 1416 | 2338.5 | 2050.5 | 112.5 | 265 | 361.49 | 33.8 | 45.4 | 0.125 | 16.75 | 0.195 | 8.5 | 0.0025 | 2.4 |
| 38 | 1416 | 2338.5 | 2050.5 | 112.5 | 265 | 361.49 | 33.8 | 45.4 | 1.25 | 167.5 | 1.95 | 85 | 0.025 | 24 |
| 39 | 1416 | 2338.5 | 2050.5 | 112.5 | 265 | 361.49 | 33.8 | 45.4 | 5 | 670 | 7.8 | 340 | 0.1 | 96 |
| 40 | 1416 | 2338.5 | 2050.5 | 112.5 | 265 | 361.49 | 33.8 | 45.4 | 5 | 670 | 7.8 | 340 | 0.1 | 480 |
| 41 | 1416 | 1559 | 1367 | 56.25 | 132.5 | 180.745 | 33.8 | 45.4 | 0.25 | 33.5 | 0.39 | 17 | 0.005 | 4.8 |
| 42 | 1416 | 1559 | 1367 | 56.25 | 132.5 | 180.745 | 16.9 | 22.7 | 0.25 | 33.5 | 0.39 | 17 | 0.005 | 4.8 |
| 43 | 1416 | 1559 | 1367 | 56.25 | 132.5 | 180.745 | 50.7 | 68.1 | 0.25 | 33.5 | 0.39 | 17 | 0.005 | 4.8 |
| 44 | 1416 | 1559 | 1367 | 56.25 | 132.5 | 180.745 | 33.8 | 45.4 | 0.125 | 16.75 | 0.195 | 8.5 | 0.0025 | 2.4 |
| 45 | 1416 | 1559 | 1367 | 56.25 | 132.5 | 180.745 | 33.8 | 45.4 | 1.25 | 167.5 | 1.95 | 85 | 0.025 | 24 |
| 46 | 1416 | 1559 | 1367 | 56.25 | 132.5 | 180.745 | 33.8 | 45.4 | 5 | 670 | 7.8 | 340 | 0.1 | 96 |
| 47 | 1416 | 1559 | 1367 | 56.25 | 132.5 | 180.745 | 33.8 | 45.4 | 5 | 670 | 7.8 | 340 | 0.1 | 480 |
| 48 | 1416 | 1559 | 1367 | 168.75 | 397.5 | 542.235 | 33.8 | 45.4 | 0.25 | 33.5 | 0.39 | 17 | 0.005 | 4.8 |
| 49 | 1416 | 1559 | 1367 | 168.75 | 397.5 | 542.235 | 16.9 | 22.7 | 0.25 | 33.5 | 0.39 | 17 | 0.005 | 4.8 |
| 50 | 1416 | 1559 | 1367 | 168.75 | 397.5 | 542.235 | 50.7 | 68.1 | 0.25 | 33.5 | 0.39 | 17 | 0.005 | 4.8 |
| 51 | 1416 | 1559 | 1367 | 168.75 | 397.5 | 542.235 | 33.8 | 45.4 | 0.125 | 16.75 | 0.195 | 8.5 | 0.0025 | 2.4 |
| 52 | 1416 | 1559 | 1367 | 168.75 | 397.5 | 542.235 | 33.8 | 45.4 | 1.25 | 167.5 | 1.95 | 85 | 0.025 | 24 |
| 53 | 1416 | 1559 | 1367 | 168.75 | 397.5 | 542.235 | 33.8 | 45.4 | 5 | 670 | 7.8 | 340 | 0.1 | 96 |
| 54 | 1416 | 1559 | 1367 | 168.75 | 397.5 | 542.235 | 33.8 | 45.4 | 5 | 670 | 7.8 | 340 | 0.1 | 480 |
| 55 | 1416 | 1559 | 1367 | 112.5 | 265 | 361.49 | 16.9 | 22.7 | 0.25 | 33.5 | 0.39 | 17 | 0.005 | 4.8 |
| 56 | 1416 | 1559 | 1367 | 112.5 | 265 | 361.49 | 16.9 | 22.7 | 0.125 | 16.75 | 0.195 | 8.5 | 0.0025 | 2.4 |
| 57 | 1416 | 1559 | 1367 | 112.5 | 265 | 361.49 | 16.9 | 22.7 | 1.25 | 167.5 | 1.95 | 85 | 0.025 | 24 |
| 58 | 1416 | 1559 | 1367 | 112.5 | 265 | 361.49 | 16.9 | 22.7 | 5 | 670 | 7.8 | 340 | 0.1 | 96 |
| 59 | 1416 | 1559 | 1367 | 112.5 | 265 | 361.49 | 16.9 | 22.7 | 5 | 670 | 7.8 | 340 | 0.1 | 480 |
| 60 | 1416 | 1559 | 1367 | 112.5 | 265 | 361.49 | 50.7 | 68.1 | 0.25 | 33.5 | 0.39 | 17 | 0.005 | 4.8 |
| 61 | 1416 | 1559 | 1367 | 112.5 | 265 | 361.49 | 50.7 | 68.1 | 0.125 | 16.75 | 0.195 | 8.5 | 0.0025 | 2.4 |
| 62 | 1416 | 1559 | 1367 | 112.5 | 265 | 361.49 | 50.7 | 68.1 | 1.25 | 167.5 | 1.95 | 85 | 0.025 | 24 |
| 63 | 1416 | 1559 | 1367 | 112.5 | 265 | 361.49 | 50.7 | 68.1 | 5 | 670 | 7.8 | 340 | 0.1 | 96 |
| 64 | 1416 | 1559 | 1367 | 112.5 | 265 | 361.49 | 50.7 | 68.1 | 5 | 670 | 7.8 | 340 | 0.1 | 480 |
| 65 | 708 | 779.5 | 683.5 | 56.25 | 132.5 | 180.745 | 33.8 | 45.4 | 0.25 | 33.5 | 0.39 | 17 | 0.005 | 4.8 |
| 66 | 708 | 779.5 | 683.5 | 112.5 | 265 | 361.49 | 16.9 | 22.7 | 0.25 | 33.5 | 0.39 | 17 | 0.005 | 4.8 |
| 67 | 708 | 779.5 | 683.5 | 112.5 | 265 | 361.49 | 33.8 | 45.4 | 0.125 | 16.75 | 0.195 | 8.5 | 0.0025 | 2.4 |
| 68 | 708 | 1559 | 1367 | 56.25 | 132.5 | 180.745 | 16.9 | 22.7 | 0.25 | 33.5 | 0.39 | 17 | 0.005 | 4.8 |
| 69 | 708 | 1559 | 1367 | 56.25 | 132.5 | 180.745 | 33.8 | 45.4 | 0.125 | 16.75 | 0.195 | 8.5 | 0.0025 | 2.4 |
| 70 | 708 | 1559 | 1367 | 112.5 | 265 | 361.49 | 16.9 | 22.7 | 0.125 | 16.75 | 0.195 | 8.5 | 0.0025 | 2.4 |
| 71 | 708 | 779.5 | 683.5 | 56.25 | 132.5 | 180.745 | 16.9 | 22.7 | 0.25 | 33.5 | 0.39 | 17 | 0.005 | 4.8 |
| 72 | 708 | 779.5 | 683.5 | 56.25 | 132.5 | 180.745 | 33.8 | 45.4 | 0.25 | 33.5 | 0.39 | 17 | 0.005 | 4.8 |
| 73 | 708 | 779.5 | 683.5 | 112.5 | 265 | 361.49 | 16.9 | 22.7 | 0.125 | 16.75 | 0.195 | 8.5 | 0.0025 | 2.4 |
| 74 | 708 | 1559 | 1367 | 56.25 | 132.5 | 180.745 | 16.9 | 22.7 | 0.125 | 16.75 | 0.195 | 8.5 | 0.0025 | 2.4 |
| 75 | 708 | 1559 | 1367 | 56.25 | 132.5 | 180.745 | 16.9 | 22.7 | 0.125 | 16.75 | 0.195 | 8.5 | 0.0025 | 2.4 |
| 76 | 2124 | 779.5 | 683.5 | 56.25 | 132.5 | 180.745 | 33.8 | 45.4 | 0.25 | 33.5 | 0.39 | 17 | 0.005 | 4.8 |
| 77 | 2124 | 779.5 | 683.5 | 112.5 | 265 | 361.49 | 16.9 | 22.7 | 0.25 | 33.5 | 0.39 | 17 | 0.005 | 4.8 |
| 78 | 2124 | 779.5 | 683.5 | 112.5 | 265 | 361.49 | 33.8 | 45.4 | 0.125 | 16.75 | 0.195 | 8.5 | 0.0025 | 2.4 |
| 79 | 2124 | 1559 | 1367 | 56.25 | 132.5 | 180.745 | 16.9 | 22.7 | 0.25 | 33.5 | 0.39 | 17 | 0.005 | 4.8 |
| 80 | 2124 | 1559 | 1367 | 56.25 | 132.5 | 180.745 | 33.8 | 45.4 | 0.125 | 16.75 | 0.195 | 8.5 | 0.0025 | 2.4 |
| 81 | 2124 | 1559 | 1367 | 112.5 | 265 | 361.49 | 16.9 | 22.7 | 0.125 | 16.75 | 0.195 | 8.5 | 0.0025 | 2.4 |
| 82 | 2124 | 779.5 | 683.5 | 56.25 | 132.5 | 180.745 | 16.9 | 22.7 | 0.25 | 33.5 | 0.39 | 17 | 0.005 | 4.8 |
| 83 | 2124 | 779.5 | 683.5 | 56.25 | 132.5 | 180.745 | 33.8 | 45.4 | 0.25 | 33.5 | 0.39 | 17 | 0.005 | 4.8 |
| 84 | 2124 | 779.5 | 683.5 | 112.5 | 265 | 361.49 | 16.9 | 22.7 | 0.125 | 16.75 | 0.195 | 8.5 | 0.0025 | 2.4 |
| 85 | 2124 | 1559 | 1367 | 56.25 | 132.5 | 180.745 | 16.9 | 22.7 | 0.125 | 16.75 | 0.195 | 8.5 | 0.0025 | 2.4 |
| 86 | 2124 | 1559 | 1367 | 56.25 | 132.5 | 180.745 | 16.9 | 22.7 | 0.125 | 16.75 | 0.195 | 8.5 | 0.0025 | 2.4 |
| 87 | 708 | 2338.5 | 2050.5 | 56.25 | 132.5 | 180.745 | 16.9 | 22.7 | 0.25 | 33.5 | 0.39 | 17 | 0.005 | 4.8 |
| 88 | 708 | 2338.5 | 2050.5 | 56.25 | 132.5 | 180.745 | 33.8 | 45.4 | 0.25 | 33.5 | 0.39 | 17 | 0.005 | 4.8 |
| 89 | 708 | 2338.5 | 2050.5 | 112.5 | 265 | 361.49 | 16.9 | 22.7 | 0.125 | 16.75 | 0.195 | 8.5 | 0.0025 | 2.4 |
| 90 | 708 | 2338.5 | 2050.5 | 56.25 | 132.5 | 180.745 | 16.9 | 22.7 | 0.125 | 16.75 | 0.195 | 8.5 | 0.0025 | 2.4 |
| 91 | 708 | 2338.5 | 2050.5 | 56.25 | 132.5 | 180.745 | 16.9 | 22.7 | 0.125 | 16.75 | 0.195 | 8.5 | 0.0025 | 2.4 |
| 92 | 708 | 779.5 | 683.5 | 56.25 | 132.5 | 180.745 | 16.9 | 22.7 | 1.25 | 167.5 | 1.95 | 85 | 0.025 | 24 |
| 93 | 708 | 779.5 | 683.5 | 56.25 | 132.5 | 180.745 | 33.8 | 45.4 | 5 | 670 | 7.8 | 340 | 0.1 | 96 |
| 94 | 708 | 779.5 | 683.5 | 112.5 | 265 | 361.49 | 16.9 | 22.7 | 5 | 670 | 7.8 | 340 | 0.1 | 480 |
| 95 | 708 | 1559 | 1367 | 56.25 | 132.5 | 180.745 | 16.9 | 22.7 | 5 | 670 | 7.8 | 340 | 0.1 | 96 |
| 96 | 708 | 1559 | 1367 | 56.25 | 132.5 | 180.745 | 16.9 | 22.7 | 5 | 670 | 7.8 | 340 | 0.1 | 480 |
| 98 | 2124 | 2338.5 | 2050.5 | 168.75 | 397.5 | 542.235 | 33.8 | 45.4 | 0.25 | 33.5 | 0.39 | 17 | 0.005 | 4.8 |
| 99 | 2124 | 2338.5 | 2050.5 | 112.5 | 265 | 361.49 | 50.7 | 68.1 | 0.25 | 33.5 | 0.39 | 17 | 0.005 | 4.8 |
| 100 | 2124 | 2338.5 | 2050.5 | 112.5 | 265 | 361.49 | 33.8 | 45.4 | 1.25 | 167.5 | 1.95 | 85 | 0.025 | 24 |
| 101 | 2124 | 2338.5 | 2050.5 | 112.5 | 265 | 361.49 | 33.8 | 45.4 | 5 | 670 | 7.8 | 340 | 0.1 | 96 |
| 102 | 2124 | 2338.5 | 2050.5 | 112.5 | 265 | 361.49 | 33.8 | 45.4 | 5 | 670 | 7.8 | 340 | 0.1 | 480 |
| 103 | 2124 | 1559 | 1367 | 168.75 | 397.5 | 542.235 | 50.7 | 68.1 | 0.25 | 33.5 | 0.39 | 17 | 0.005 | 4.8 |
| 104 | 2124 | 2338.5 | 2050.5 | 168.75 | 397.5 | 542.235 | 33.8 | 45.4 | 1.25 | 167.5 | 1.95 | 85 | 0.025 | 24 |
| 105 | 2124 | 2338.5 | 2050.5 | 168.75 | 397.5 | 542.235 | 33.8 | 45.4 | 5 | 670 | 7.8 | 340 | 0.1 | 96 |
| 106 | 2124 | 2338.5 | 2050.5 | 168.75 | 397.5 | 542.235 | 33.8 | 45.4 | 5 | 670 | 7.8 | 340 | 0.1 | 480 |
| 107 | 2124 | 2338.5 | 2050.5 | 112.5 | 265 | 361.49 | 50.7 | 68.1 | 1.25 | 167.5 | 1.95 | 85 | 0.025 | 24 |

| | | | | | | | | | | | | | | |
|---|---|---|---|---|---|---|---|---|---|---|---|---|---|---|
| 108 | 2124 | 2338.5 | 2050.5 | 112.5 | 265 | 361.49 | 50.7 | 68.1 | 5 | 670 | 7.8 | 340 | 0.1 | 96 |
| 109 | 2124 | 2338.5 | 2050.5 | 112.5 | 265 | 361.49 | 50.7 | 68.1 | 5 | 670 | 7.8 | 340 | 0.1 | 480 |
| 110 | 2124 | 1559 | 1367 | 168.75 | 397.5 | 542.235 | 50.7 | 68.1 | 0.25 | 33.5 | 0.39 | 17 | 0.005 | 4.8 |
| 111 | 2124 | 1559 | 1367 | 168.75 | 397.5 | 542.235 | 33.8 | 45.4 | 1.25 | 167.5 | 1.95 | 85 | 0.025 | 24 |
| 112 | 2124 | 1559 | 1367 | 168.75 | 397.5 | 542.235 | 33.8 | 45.4 | 5 | 670 | 7.8 | 340 | 0.1 | 96 |
| 113 | 2124 | 2338.5 | 2050.5 | 168.75 | 397.5 | 542.235 | 33.8 | 45.4 | 5 | 670 | 7.8 | 340 | 0.1 | 480 |
| 114 | 2124 | 2338.5 | 2050.5 | 112.5 | 265 | 361.49 | 50.7 | 68.1 | 1.25 | 167.5 | 1.95 | 85 | 0.025 | 24 |
| 115 | 2124 | 2338.5 | 2050.5 | 112.5 | 265 | 361.49 | 50.7 | 68.1 | 5 | 670 | 7.8 | 340 | 0.1 | 96 |
| 116 | 2124 | 2338.5 | 2050.5 | 112.5 | 265 | 361.49 | 50.7 | 68.1 | 5 | 670 | 7.8 | 340 | 0.1 | 480 |
| 117 | 2124 | 1559 | 1367 | 112.5 | 265 | 361.49 | 50.7 | 68.1 | 1.25 | 167.5 | 1.95 | 85 | 0.025 | 24 |
| 118 | 2124 | 1559 | 1367 | 112.5 | 265 | 361.49 | 50.7 | 68.1 | 5 | 670 | 7.8 | 340 | 0.1 | 96 |
| 119 | 2124 | 1559 | 1367 | 112.5 | 265 | 361.49 | 50.7 | 68.1 | 5 | 670 | 7.8 | 340 | 0.1 | 480 |
| 120 | 2124 | 2338.5 | 2050.5 | 168.75 | 397.5 | 542.235 | 50.7 | 68.1 | 1.25 | 167.5 | 1.95 | 85 | 0.025 | 24 |
| 121 | 2124 | 2338.5 | 2050.5 | 168.75 | 397.5 | 542.235 | 50.7 | 68.1 | 5 | 670 | 7.8 | 340 | 0.1 | 96 |
| 122 | 2124 | 2338.5 | 2050.5 | 168.75 | 397.5 | 542.235 | 50.7 | 68.1 | 5 | 670 | 7.8 | 340 | 0.1 | 480 |

S2. Salt components included for Validation 1 treatments. All values represent mg/L. Additional components of these treatments included 100mg/L Myo-Inositol, 2mg/L Glycine, 1mg/L Nicotinic acid, 2mg/L Thiamine, 0.004mg/L CoCl2, and 0.1mg/L KI, which were added consistently to all treatments. Additionally, each treatment included 0.6% (w/v) agar, 3% (w/v) sucrose, and pH = 5.7.

| Treatment | NH4NO3 | K2SO4 | Ca(NO3)2 | CaCl2 | KH2PO4 | MgSO4 | FeSO4 · 7H2O | Na2EDTA · 2H2O | CuSO4 · 5H2O | MnSO4 · H2O | Na2MoO4 · 2H2O | Zn(NO3)2.6H2O | NiSO4.6H2O | H3BO3 |
|---|---|---|---|---|---|---|---|---|---|---|---|---|---|---|
| 1 | 1862.479 | 1531.472 | 767.730 | 91.123 | 364.441 | 343.465 | 28.668 | 26.861 | 3.253 | 342.795 | 0.393 | 259.889 | 0.073 | 3.403 |
| 2 | 2053.514 | 1683.259 | 1089.056 | 68.199 | 196.287 | 372.459 | 48.277 | 27.422 | 2.487 | 297.227 | 5.219 | 23.141 | 0.036 | 138.492 |
| 3 | 1435.301 | 781.883 | 692.961 | 116.879 | 273.280 | 368.915 | 37.867 | 40.185 | 0.178 | 18.475 | 0.200 | 9.116 | 0.003 | 2.370 |
| 4 | 1419.521 | 781.708 | 679.328 | 101.205 | 269.947 | 368.366 | 78.398 | 63.154 | 0.358 | 36.115 | 0.460 | 18.088 | 0.081 | 4.714 |
| 5 | 1741.500 | 1408.285 | 1517.541 | 60.748 | 349.214 | 493.818 | 46.220 | 54.006 | 4.688 | 31.827 | 7.117 | 53.772 | 0.076 | 189.496 |
| 6 | 1897.586 | 1754.129 | 1148.641 | 140.262 | 389.076 | 378.113 | 24.269 | 28.248 | 0.156 | 140.529 | 2.946 | 57.434 | 0.013 | 293.804 |
| 7 | 2090.636 | 2073.124 | 1057.746 | 71.688 | 172.599 | 388.630 | 35.168 | 44.742 | 3.947 | 260.789 | 2.204 | 41.251 | 0.046 | 265.592 |
| 8 | 1485.925 | 2337.944 | 1953.434 | 110.907 | 273.280 | 368.915 | 34.117 | 39.685 | 0.128 | 17.225 | 0.224 | 8.650 | 0.003 | 2.649 |
| 9 | 1533.757 | 787.055 | 1114.461 | 68.528 | 137.228 | 396.657 | 46.209 | 58.070 | 3.379 | 55.225 | 3.087 | 39.650 | 0.055 | 73.891 |
| 10 | 1355.886 | 881.693 | 771.301 | 58.733 | 134.697 | 443.284 | 41.901 | 59.797 | 1.918 | 42.895 | 2.220 | 130.407 | 0.026 | 18.320 |
| 11 | 2067.337 | 1065.058 | 848.196 | 110.907 | 173.280 | 419.040 | 47.117 | 39.685 | 0.628 | 131.620 | 4.824 | 133.088 | 0.045 | 91.281 |
| 12 | 1418.363 | 1560.736 | 1374.462 | 173.736 | 393.580 | 546.482 | 37.398 | 50.239 | 0.276 | 31.403 | 0.419 | 19.484 | 0.006 | 5.967 |
| 13 | 785.925 | 1588.613 | 1408.890 | 54.744 | 137.635 | 188.915 | 17.117 | 23.685 | 0.328 | 32.225 | 0.387 | 16.650 | 0.005 | 4.249 |
| 14 | 2067.337 | 781.948 | 689.255 | 110.870 | 271.256 | 342.273 | 16.222 | 22.179 | 0.311 | 34.863 | 0.373 | 16.963 | 0.006 | 5.281 |
| 15 | 2119.301 | 2137.883 | 2049.461 | 112.879 | 269.112 | 368.915 | 47.117 | 69.685 | 1.228 | 166.225 | 1.956 | 85.116 | 0.027 | 23.970 |
| PHJ | 2067.337 | 2337.948 | 2050.133 | 161.446 | 394.256 | 542.273 | 50.222 | 68.179 | 1.096 | 167.620 | 1.824 | 86.963 | 0.029 | 26.281 |

S3. Salt components included for Validation 2 treatments. All values represent mg/L. Additional components of these treatments included 100mg/L Myo-Inositol, 2mg/L Glycine, 1mg/L Nicotinic acid, 2mg/L Thiamine, 0.004mg/L CoCl2, and 0.1mg/L KI, which were added consistently to all treatments. Additionally, each treatment included 0.6% (w/v) agar, 3% (w/v) sucrose, and pH = 5.7.

| Treatment | NH4NO3 | K2SO4 | Ca(NO3)2 | CaCl2 | KH2PO4 | MgSO4 | FeSO4 · 7H2O | Na2EDTA · 2H2O | CuSO4 · 5H2O | MnSO4 · H2O | Na2MoO4 · 2H2O | Zn(NO3)2.6H2O | NiSO4.6H2O | H3BO3 |
|---|---|---|---|---|---|---|---|---|---|---|---|---|---|---|
| 1 | 1533.757 | 787.055 | 1114.461 | 68.528 | 137.228 | 396.657 | 46.209 | 58.070 | 3.379 | 55.225 | 3.087 | 39.650 | 0.055 | 73.891 |
| 2 | 1355.886 | 881.693 | 771.301 | 58.733 | 134.697 | 443.284 | 41.901 | 59.797 | 1.918 | 42.895 | 2.220 | 130.407 | 0.026 | 18.320 |
| 3 | 1418.363 | 1560.736 | 1374.462 | 163.736 | 393.580 | 541.482 | 37.398 | 50.239 | 0.276 | 31.403 | 0.419 | 19.484 | 0.006 | 5.967 |
| 4 | 785.925 | 1588.613 | 1408.890 | 56.744 | 137.635 | 188.915 | 17.117 | 23.685 | 0.328 | 32.225 | 0.387 | 16.650 | 0.005 | 4.249 |
| 5 | 2119.301 | 2137.883 | 2049.461 | 112.879 | 269.112 | 368.915 | 47.117 | 67.685 | 1.228 | 166.225 | 1.956 | 85.116 | 0.027 | 23.970 |
| PHJ | 2067.337 | 2337.948 | 2050.133 | 161.446 | 394.256 | 542.213 | 50.222 | 68.079 | 1.096 | 167.620 | 1.824 | 86.963 | 0.029 | 26.281 |
| PHJ-1 | 2123.239 | 1557.239 | 1363.220 | 167.225 | 286.419 | 301.087 | 27.918 | 38.650 | 0.127 | 17.026 | 0.197 | 9.419 | 0.004 | 6.228 |

S4. Effect of PHJ and PHJ+PA on different explant types, including shoot tip (ST) and nodal cutting (NC), following the subculture 1 phase. Values represent averages of the parameters tested ± standard error of the mean. Significance, with 95% confidence, is noted with asterisks.

| Parameter | Media | ST | NC |
|---|---|---|---|
| Shoot Number | PHJ | 1.84 ± 0.12 | 1.91 ± 0.133 |
| | PHJ+PA | 2.72 ± 0.60 | 4.45 ± 0.44** |
| Number of Nodes | PHJ | 8.84 ± 0.46 | 8.25 ± 0.47 |
| | PHJ+PA | 8.25 ± 0.91 | 9.45 ± 0.63* |
| Canopy Area | PHJ | 1564.98 ± 106.24 | 3120.65 ± 320.27 |
| | PHJ+PA | 2414.2 ± 632.42 | 3685.34 ± 573.17 |

S5. Effect of DKW and DKW+PA on different explant types, including shoot tip (ST) and nodal cutting (NC), following the subculture 1 phase. Values represent averages of the parameters tested ± standard error of the mean. Significance, with 95% confidence, is noted with asterisks.

| Parameter | Media | ST | NC |
|---|---|---|---|
| Shoot Number | DKW | 1.25 ± 0.09 | 1.6875 ± 0.15 |
| | DKW+PA | 1.90625 ± 0.3* | 2.78125 ± 0.63* |
| Number of Nodes | DKW | 6.71875 ± 0.21 | 5.8125 ± 0.31 |
| | DKW+PA | 6.78125 ± 0.79 | 7.28125 ± 0.88 |
| | DKW | 1805.695 ± 155.24 | 1463.778 ± 139.82 |

| | | | |
|---|---|---|---|
| Canopy Area | DKW+PA | 1451.636 ± 436.98 | 3175.344 ± 1024.11 |

S6. Effect of PHJ and PHJ+PA on different explant types, including shoot tip (ST) and nodal cutting (NC), following the subculture 2 phase. Values represent averages of the parameters tested ± standard error of the mean. Significance, with 95% confidence, is noted with asterisks.

| Parameter | Media | ST | NC |
|---|---|---|---|
| Shoot Number | PHJ | 1.66 ± 0.29 | 2.25 ± 0.16 |
| | PHJ+PA | 4.78 ± 1.15* | 5.11 ± 0.38*** |
| Number of Nodes | PHJ | 9.5 ± 0.36 | 9.41 ± 0.43 |
| | PHJ+PA | 9.84 ± 0.94 | 11.25 ± 0.57* |
| Canopy Area | PHJ | 2082.17 ± 278.52 | 3903.4 ± 608.04 |
| | PHJ+PA | 5648.17 ± 1678.346* | 7009.35 ± 1342.46 |

S7. Effect of DKW and DKW+PA on different explant types, including shoot tip (ST) and nodal cutting (NC), following the subculture 2 phase. Values represent averages of the parameters tested ± standard error of the mean. Significance, with 95% confidence, is noted with asterisks.

| Parameter | Media | ST | NC |
|---|---|---|---|
| Shoot Number | DKW | 1.28 ± 0.1 | 1.78 ± 0.14 |
| | DKW+PA | 2.03 ± 0.17*** | 3.53 ± 0.79* |
| Number of Nodes | DKW | 8.5 ± 0.34 | 7.66 ± 0.58 |
| | DKW+PA | 8.44 ± 0.54 | 8.8125 ± 1.15 |
| Canopy Area | DKW | 2991.43 ± 620.41 | 3781.56 ± 658.66 |
| | DKW+PA | 2894.43 ± 734.93 | 3991.92 ± 1185.69 |

S8. Effect of PHJ and PHJ+PA on different explant types, including shoot tip (ST) and nodal cutting (NC), following the subculture 3 phase. Values represent averages of the parameters tested ± standard error of the mean. Significance, with 95% confidence, is noted with asterisks.

| Parameter | Media | ST | NC |
|---|---|---|---|
| Shoot Number | PHJ | 1.28 ± 0.14 | 2.25 ± 0.18 |
| | PHJ+PA | 2.5 ± 0.34*** | 3.34 ± 0.39** |
| Number of Nodes | PHJ | 8.38 ± 0.37 | 7.25 ± 0.3 |
| | PHJ+PA | 9.03 ± 0.41 | 9.22 ± 0.5** |
| Canopy Area | PHJ | 2222.48 ± 113.77 | 1807.15 ± 150.26 |
| | PHJ+PA | 3596.95 ± 1009.14 | 2961.91 ± 410.78* |

S9. Effect of DKW and DKW+PA on different explant types, including shoot tip (ST) and nodal cutting (NC), following the subculture 3 phase. Values represent averages of the parameters tested ± standard error of the mean. Significance, with 95% confidence, is noted with asterisks.

| Parameter | Media | ST | NC |
|---|---|---|---|
| Shoot Number | DKW | 1.25 ± 0.7 | 1.63 ± 0.12 |
| | DKW+PA | 1.34 ± 0.15 | 1.88 ± 0.28 |
| Number of Nodes | DKW | 7.16 ± 0.62* | 5.59 ± 0.22 |
| | DKW+PA | 6.0313 ± 0.55 | 6.47 ± 0.8 |
| Canopy Area | DKW | 2118.44 ± 229.47 | 1712.79 ± 87.28 |
| | DKW+PA | 1468.18 ± 481.18 | 2221.5 ± 519.45 |

S10. Assessment of SLH plantlets on PHJ and DKW following three rounds of subculture after culture initiation. Values represent averages of the parameters tested ± standard error of the mean. Significance, with 95% confidence, is noted with asterisks.

| Parameter | Media | SLH |
|---|---|---|
| Shoot Number | PHJ | 1.63 ± 0.16* |
| | DKW | 1 |
| Number of Nodes | PHJ | 5.63 ± 0.6* |
| | DKW | 3.13 ± 0.22 |

S11. Assessment of SLH plantlets on PHJ+PA and DKW+PA following three rounds of subculture after culture initiation. Values represent averages of the parameters tested ± standard error of the mean. Significance, with 95% confidence, is noted with asterisks.

| Parameter | Media | SLH | TRS10 |
|---|---|---|---|
| Shoot Number | PHJ+PA | 2.81 ± 0.31 | 3.5 ± 0.74 |
| | DKW+PA | 2.56 ± 0.44 | 2.56 ± 0.21 |
| Number of Nodes | PHJ+PA | 6.06 ± 0.43 | 8.88 ± 0.13** |
| | DKW+PA | 5.25 ± 0.27 | 5.06 ± 0.37 |
